# Muscle Stiffness, Not Force, Limits Human Force Production

**DOI:** 10.1101/2025.09.03.673119

**Authors:** F. Tessari, N. Hogan

**Affiliations:** Department of Mechanical Engineering, Massachusetts Institute of Technology, Cambridge, USA; Department of Brain and Cognitive Sciences, Massachusetts Institute of Technology, Cambridge, USA

## Abstract

Humans have a remarkable ability to manage physical interaction despite a tremendously complex musculoskeletal system, comparatively slow muscles and slow neural communication. Most physical interaction tasks require force generation, and that can compromise stability. To ensure stability, muscles are required to produce a certain mechanical impedance (especially stiffness and damping). In this work we investigate the human limits of force generation and test whether and how those limits are constrained by our ability to stabilize the task. Moreover, we also study how strongly limb configuration – known to play a key role in transmitting muscle tension to hand force – matters in the generation of force. We devised an experiment in which healthy individuals performed a maximum voluntary pushing task using a custom-designed gimbal apparatus which allowed us to conditionally change the ability to transmit torque at the wrist, thus enabling or disabling stabilization via wrist pronation-supination, ulnar-radial deviation, and flexion-extension. The experiment was repeated in two different arm configurations (preferred and undesired) to assess the effect of limb configuration. The results showed: (i) a statistically significant and substantial decrease in maximum output force with less wrist ability to generate stabilizing torque, and (ii) a significant but minimal difference in maximum pushing force between the two arm configurations. Interestingly, subjects unexpectedly improved their maximum pushing performance between pre- and post-experiment, suggesting a negligible effect of fatigue and a possible effect of learning. Our findings confirm our hypothesis that force production appears to be limited by stiffness production. Muscles do not primarily generate force, but generate mechanical impedance.

## Introduction

Humans are extremely good at managing physical interaction despite: (i) a complex musculoskeletal system with more than 200 degrees of freedom and 600+ muscles, (ii) slow neural communication (∼10^2^ m/s), (iii) substantial transmission delays (∼100-200 ms), and (iv) limited muscle control bandwidth (≪ 10 Hz) (Hogan, 2017; Kandel et al., 2021).

Most physical interaction tasks require the production of force. Imagine, for example, hammering a nail, opening a bottle, or holding a mug in your hand. However, we often overlook the fact that force production can compromise the mechanical stability of some tasks. Two simple examples are pushing a screwdriver or bench pressing with free weights. In both cases, the generation of force also produces a destabilizing action which has the tendency to push the limbs away from their nominal posture. Small deviation around the direction of force generation – when pushing a screw driver or lifting a dumbbell – will generate a de-stabilizing force which – if not compensated – will tend to increase the deviation, leading to catastrophic failure. Figure 1 provides an intuitive representation of the problem.

**Figure 1.**
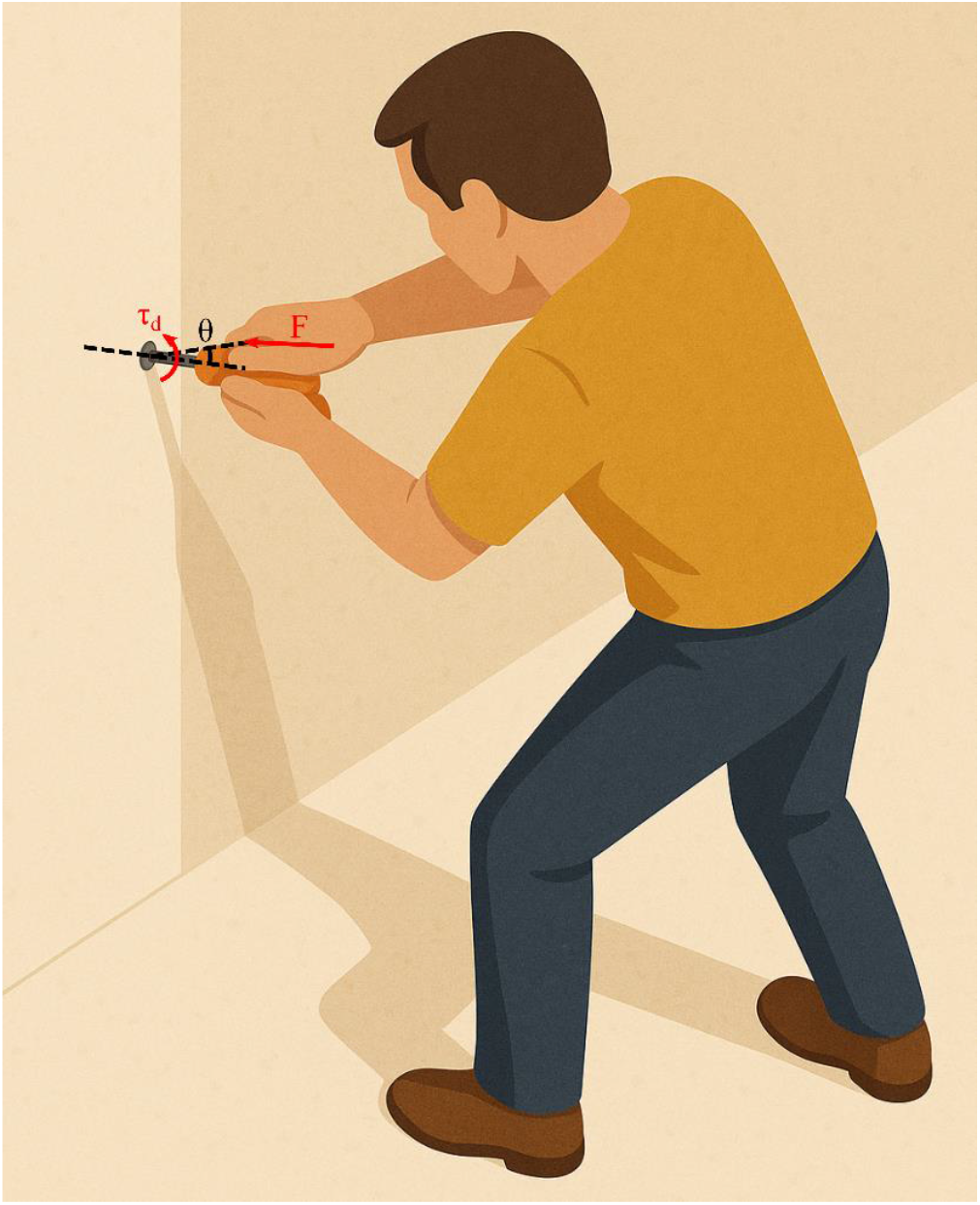
Illustration exemplifying the destabilizing nature of some physical interaction tasks. In the depicted example, an individual is applying a certain force F to a screw-driver against a screw mounted on a vertical wall. The screw-driver is not perfectly aligned. by an angle θ. with the normal direction to the wall. This means that force F will also generate a destabilizing action (τ_d_), that. if not compensated. will lead to ‘catastrophic failure’ of the task. [Illustration background generated using AI support. Mathematical notation added by the authors.]

Humans can compensate for the destabilizing effect of force production by generating mechanical impedance, and specifically stiffness (Hogan, 1985). Several studies have investigated the ability of humans to generate and regulate impedance^1^, and particularly stiffness, during physical interaction (Bennett et al., 1992; Franklin et al., 2003; Gomi & Kawato, 1997; Höppner et al., 2017; Kennedy & Schwartz, 2019; Loram & Lakie, 2002; Ludvig & Kearney, 2007; Nichols & Houk, 1976; Perreault et al., 2001, 2002; Phan et al., 2020; Takagi et al., 2020; Zhang et al., 2025). Existing experimental results have quantified the human ability to tune muscle, joint, and end-effector impedance to maintain stability during physical interaction in both perturbed (Burdet et al., 2001; Perreault et al., 2001) and unperturbed (Zhang et al., 2025) conditions. In a previous work from our group (Zhang et al., 2025), we showed how healthy individuals are capable of predictively regulating their arm end-effector stiffness to succeed in a challenging physical interaction task. Moreover, we also observed that subjects tended to use the minimum allowable stiffness to do so.

Note, however, that not all physical interaction tasks are intrinsically destabilizing. Simple examples include holding a mug of coffee, or pushing against a flat wall. Two questions arise: (i) Is the human ability to generate force limited by the ability to stabilize posture? In other words, which is the limiting factor in force generation? (ii) How critical is the posture chosen to perform a force-exertion task? We know that arm pose and joint configuration play a key role in transmitting muscle tension to hand force. Moreover, humans appear to exhibit a preference for certain poses over others. For example, most upper-limb manipulation tasks are preferentially performed about a half meter below the shoulder and about a half meter in front of the sternum. This suggests a “motor fovea” (by analogy with the visual fovea (Schira et al., 2009)) where dexterity and manipulability are maximized. However, we also know that the redundancy of the human musculoskeletal system allows for multiple solutions to emerge for the same motor task (Bernstein, 1967; Latash, 2012; Tessari et al., 2025). Does deviation from a preferred posture affect the ability to recruit muscles and limit the ability to exert force?

To address these questions, we designed an experiment in which healthy individuals were tasked to perform a simple pushing task with maximum effort. They pushed on an experimental apparatus that resembled a hand tool but included a lockable gimbal device. With it, we tested different levels of inhibition of the wrist’s ability to generate stabilizing torques in both preferred and undesired arm configurations.

Our results showed that (i) human maximum force production – during a pushing task – is limited by the ability to generate stiffness, and that (ii) substantial deviation of the arm configuration from its preferred posture had a negligible effect on performance. Interestingly, we also found that, with exposure, healthy individuals improved their performance of this very familiar and common activity, pushing.

## Materials & Methods

### Stabilizing an Unstable Pushing Task

Several tasks involving force exertion on a curved surface can lead to static instability. Consider for example the common tasks of pushing a stick, a screw driver or a powered drill, against a vertical wall (refer to Figure 1). As long as the stick is perfectly perpendicular (*θ* = 90^°^) to the wall, and the force *F* is applied along the perpendicular direction, the system is in static equilibrium. However, if a small perturbation appears – for example due to the presence of noise in the force generator e.g., the human musculoskeletal system, or noise in the environment e.g., a gust of wind or a vibration – the resulting angular deviation (Δ*θ*) from the perpendicular direction will generate a destabilizing moment *τ*_*d*_, which –if not compensated – will lead to a catastrophic failure of the task. Figure 2 provides a schematic illustration of the mechanics of the task.

**Figure 2.**
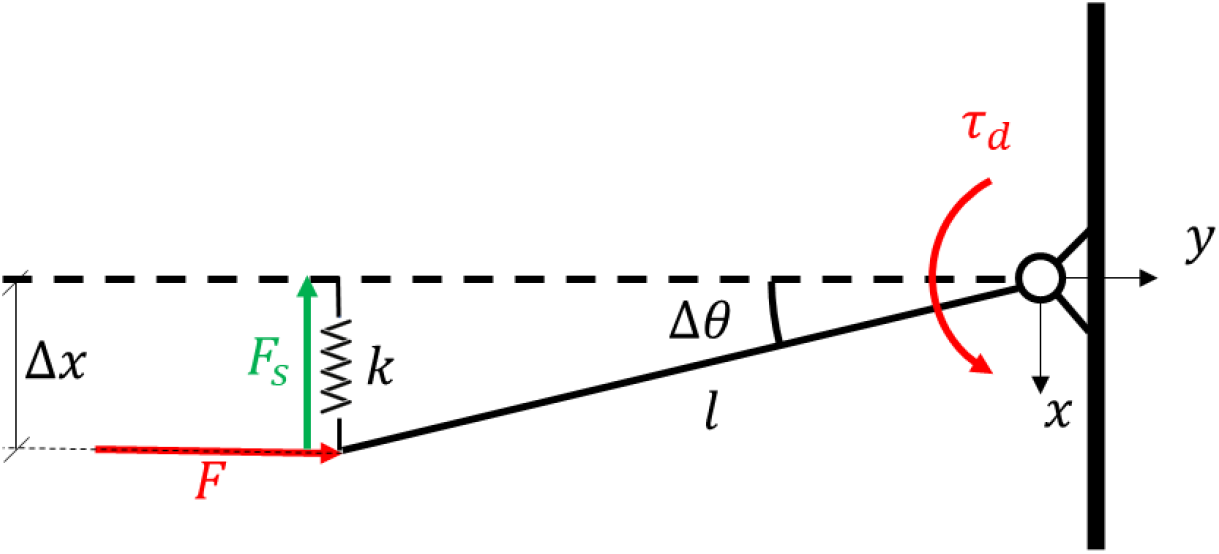
Schematic illustration of the destabilizing action of pushing on a stick. The stick of length ‘l’ is modeled as a beam hinged at the wall. A force ‘F’ (red) is applied along the ‘y’ direction, perpendicular to the wall. A small perturbation causes a deviation ‘Δx’ at the tip of the stick which corresponds to an angular deviation ‘Δ*θ*’. This deviation causes the generation of a destabilizing moment ′τ_d_′. A force ‘F_s_’ (green) is required to stabilize the task. A spring of sufficient stiffness ‘k’ can generate the required stabilizing force.

In order to compensate for such a moment, the user needs to introduce a stabilizing force *F*_*s*_ which can be estimated knowing the length of the stick and the deviation from the perpendicular direction.

In mathematical terms, the destabilizing torque is equal to:

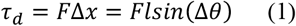

which if we assume small angular deviations, Δ*θ* ≈ 0 → sin(Δ*θ*) ≈ Δ*θ*, cos(Δ*θ*) ≈ 1, returns:

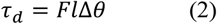

The stabilizing force needs to generate a moment of equal or greater magnitude and opposite sign of *τ*_*d*_. This results in:

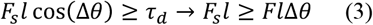

A reasonable way to passively generate the required stabilizing force ‘*F*_*s*_’ is by imposing a force proportional to the observed displacement i.e., introducing a stiffness (vertical in the specific case of Figure 2):

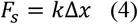

Combining Equations (4) and (3), we obtain:

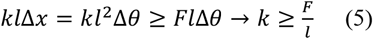

The obtained value of ‘*k*’ represents the minimum required translational stiffness to guarantee stability in an unstable pushing task. This translational stiffness can be generated by a suitable rotational stiffness about the joints e.g., the wrist and/or the shoulder. For example, consider a planar arm model – as depicted in Figure 3 – and assume that: (i) the elbow is generating a pushing force that is aligned with respect to the wrist-shoulder line, and (ii) the deviations are small such that Δ*x* ≈ *l*Δ*θ* ≈ *r*Δ*α*.

**Figure 3.**
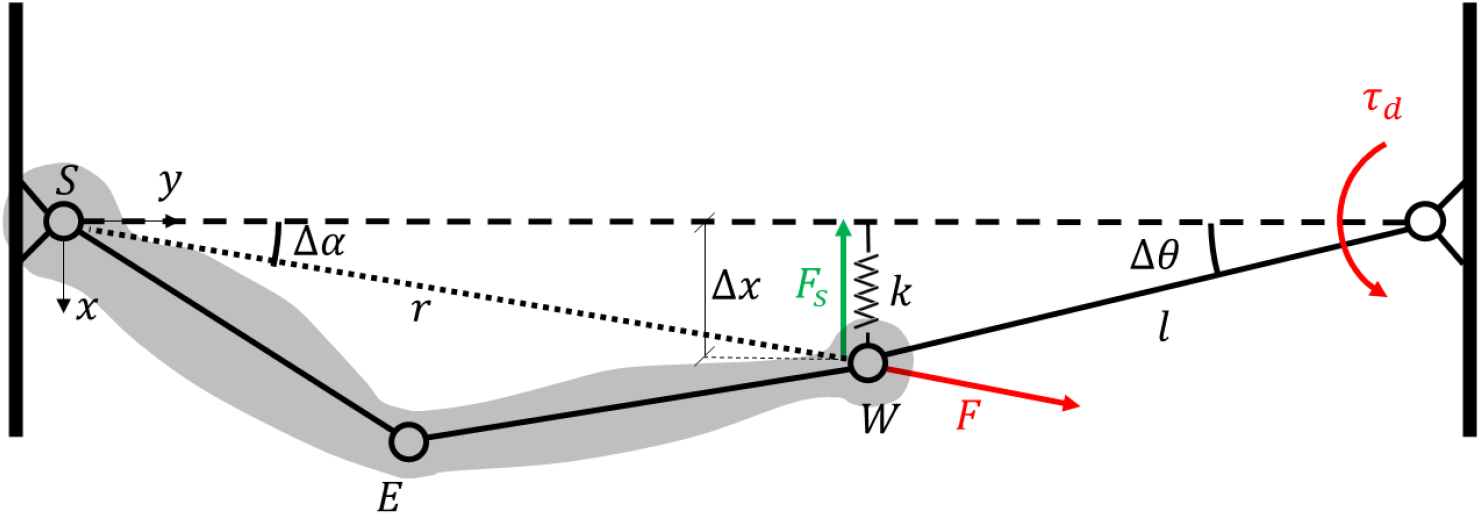
Schematic illustration of the destabilizing action of pushing on a stick with a coupled planar arm model. The stick of length ‘l’ is modeled as a beam hinged at the wall. The arm is modeled as 3 DoF planar manipulandum. The pushing force ‘F’ (red) is applied along the shoulder-wrist line of length ‘r’, which deviates by angle ‘Δα’ from the ‘y’ direction. A small perturbation causes a deviation ‘Δx’ at the tip of the stick which corresponds to an angular deviation ‘Δ*θ*’. This deviation causes the generation of a destabilizing moment ′τ_d_′. A force ‘F_s_’ (green) is required to stabilize the task. A spring of sufficient stiffness ‘k’ can generate the required stabilizing force.

In this scenario, the minimum required translational stiffness becomes^2^:

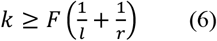

Interestingly, in both analyzed cases the minimum required translational stiffness (see Equations 5 and 6) increases with the amount of force we need to generate, and decreases with both the length of the stick ‘*r*’ and the wrist-shoulder distance ‘*l*’.

We can now map the minimum required translational stiffness to either the shoulder or wrist joint. To do so, the shoulder or the wrist have to generate a torque that compensates for the destabilizing action of the force ‘*F*’. The shoulder torque can be modeled as *τ*_*s*_ = *k*_*s*_Δα, while wrist torque can be modeled as *τ*_*w*_ = *k*_*w*_(Δα + Δ*θ*). ‘*k*_*s*_’ represents the minimum required shoulder joint stiffness (acting alone), and ‘*k*_*w*_’ represents the minimum required wrist joint stiffness (acting alone). By equating the shoulder and wrist torques to the stabilizing action of ‘*F*_*s*_’ or the destabilizing action of ‘*F*’, we obtain:

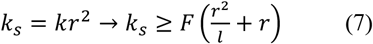

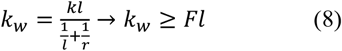

If we observe how the translation stiffness maps to the wrist or shoulder (Equations 7 and 8), we notice that the minimum wrist and shoulder stiffness increase with the amount of generated force ‘*F*’. Moreover, the wrist stiffness also grows with length of the stick ‘*l*’, while the shoulder stiffness decreases with it. We invite readers to review the preliminary works by Rancourt and Hogan for a more detailed mathematical description of the biomechanics of force production (Rancourt & Hogan, 2001, 2009).

The important methodological result from Equations 5,6,7, and 8, is that – in all cases – force and stiffness are intrinsically connected in destabilizing physical interaction tasks.

### Experimental Setup

Nine healthy individuals, (5F,4M), age 24 ± 4 years old, were recruited to perform the experiment. All subjects but one were right-handed; subject number 5 was ambidextrous. All subjects performed the experiment with the right hand. They all received a comprehensive briefing on the experimental procedures, and signed a consent form approved by MIT’s Institutional Review Board (IRB protocol nr. 2408001378).

Subjects were instructed to sit on a stool with adjustable height and no back support. The height of the stool was adjusted such that the subject could comfortably maintain both feet in contact with the ground at all times. Subjects were instructed to maintain an upright posture. Nonetheless, leaning back or forth during the experiment was allowed. The stool was positioned in front of a Kuka LBR iiwa robot equipped at its end-effector with an ATI 6-axis force-torque sensor (US-30-100 Gamma series). The sensor was used to measure the interaction forces and torques of the subjects during the task. The decision to mount the sensor on a programmable robot was motivated by the need to adjust the placement of the sensor with respect to the subject.

For each subject the robot was programmed to maintain its end-effector at a horizontal distance (‘z’ direction in Figure 3) of 43 ± 2 *cm* from the subjects’ shoulder, and at a height of 98 ± 3 *cm* from the ground (‘x’ direction in Figure 4). The robot end-effector was then aligned in the medial direction (‘y’ direction in Figure 4) such that the subjects’ hand – when in contact with the robot – would remain to the right of the right shoulder^3^. The exact location was adjusted, on a subject-by-subject basis, until each subject reported that it was his/her preferred configuration to perform the task i.e., pushing. Please refer to Figure 4 for a visual representation of the experimental setup.

**Figure 4.**
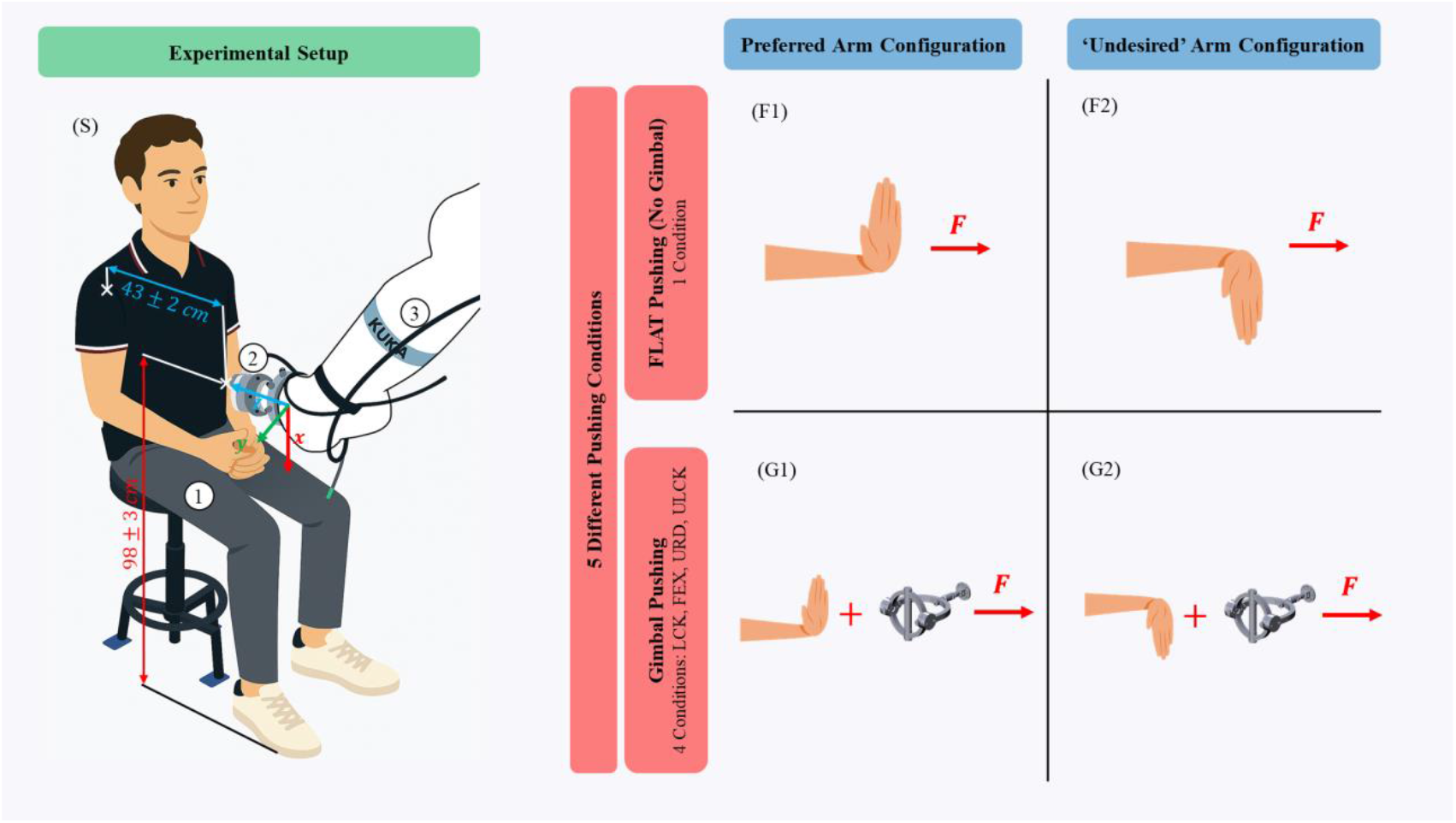
Panel (S) presents an illustration of the Experimental Setup. The subject (1) was seated on an adjustable stool in front of a Kuka LBR iiwa (3) with a 6-axis ATI force/torque sensor (2) mounted at its end-effector. The sensor was placed at a height of 98 ± 3 cm from the ground (‘x’ axis, red). The subject’s shoulder center was located at distance of 43 ± ^2^ cm from the force sensor along the ‘z’ axis (blue). The lateral position (‘y’ axis, green) was adjusted on a subject-by-subject basis to maximize pushing comfort. The four panels on the right represent the different experimental conditions. Specifically, the experiment was organized in 4+1 different pushing conditions (‘FLAT’ pushing without the gimbal device, and ‘LCK’, ‘FEX’, ‘URD’, ‘ULCK’ with the gimbal device) and 2 different arm configurations (‘preferred’ and ‘undesired’). The ‘FLAT’ pushing conditions for the ‘preferred’ and ‘undesired’ arm configuration are reported in panels (F1) and (F2), respectively. The gimbal pushing conditions for the ‘preferred’ and ‘undesired’ arm configuration are reported in panels (G1) and (G2), respectively. In the gimbal cases, the subjects were instructed to grasp the handle of the gimbal to perform the pushing task.

A custom gimbal device was designed and manufactured to progressively modify the ability of the wrist to generate torque. Figure 5 provides a detailed view of the apparatus. It consists of three axes of rotation, mutually orthogonal and intersecting at one point. This allowed the gimbal to reproduce the behavior of a spherical joint. Each axis of rotation could be locked by means of mechanical pins which could be manually inserted in the system. When an axis of rotation was unlocked, the wrist was unable to transmit torque in that direction, and vice versa. The gimbal was connected to the robot by means of a ball joint. This was implemented to closely reproduce the common task of pushing on a drill or screwdriver against a wall – similar to what is illustrated in Figures 1 and 2. The distance between the ball joint and the intersection of the 3 axes was 238.44 mm. The gimbal device CAD model and related bill of materials are freely available online at https://github.com/ftessari23/GimbalPaper.

**Figure 5.**
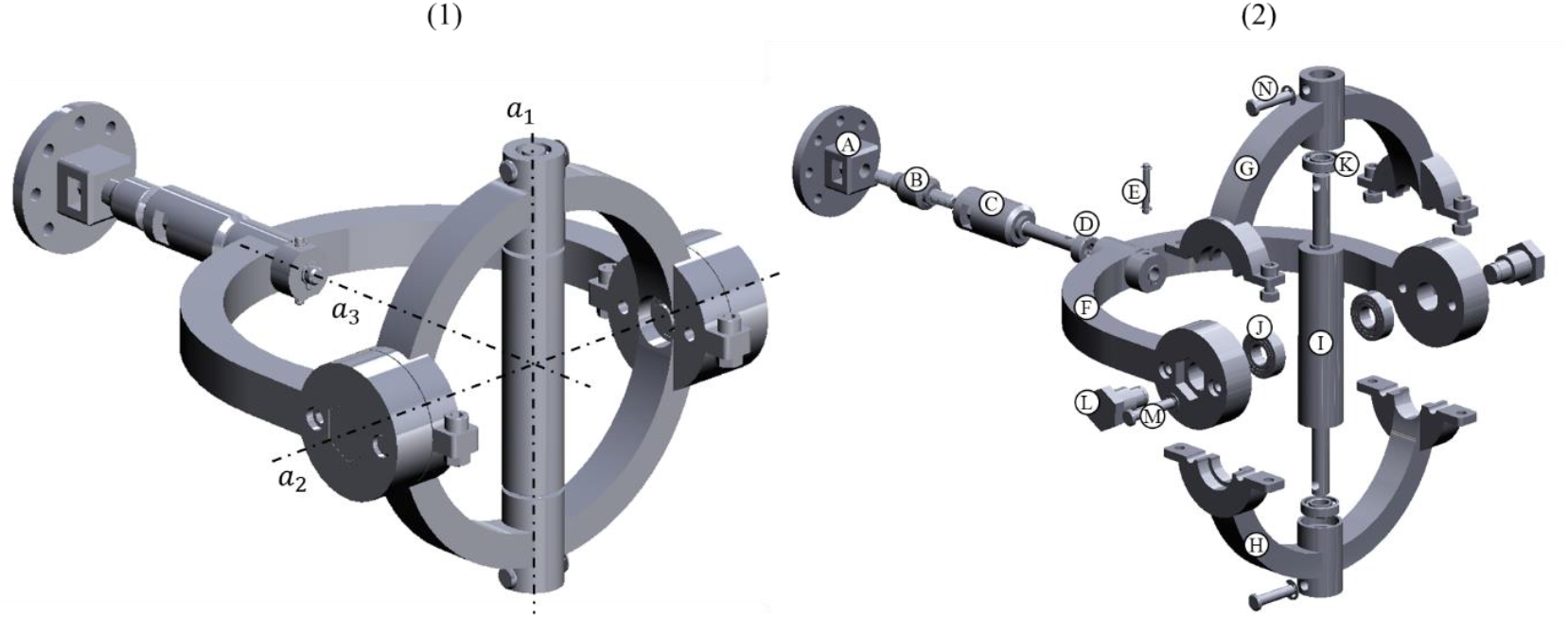
Panel (1) presents the 3D CAD model of the gimbal device. The three lockable axes of rotation are reported with dot-dashed black lines: ‘a_1_’ flexion-extension, ‘a^2^’ ulnar-radial deviation, and ‘a_3_’ pronation-supination. Panel (2) shows an exploded view of the 3D CAD models: (A) connection plate to ATI force/torque sensor, (B) ball joint, (C) stick, (D) thrust bearing, (E) pin for locking axis 3, (F) fork, (G), top orbit, (H) bottom orbit, (I) roller, (J) ball bearing for axis 2, (K) ball bearing for axis 1, (L) connection pin, (M) pin for locking axis 2, (N) pin for locking axis 1. The CAD model and related bill-of-materials are freely available at https://github.com/ftessari23/GimbalPaper.

### Experimental Protocol

Before initiating the experiment, subjects completed a brief survey reporting their age, weight, height and handedness. The length of the upper arm (acromion to olecranon), forearm (olecranon to styloid process), and hand (styloid process to middle finger knuckle) were manually measured using a measuring tape.

After completing the survey, subjects were placed in front of the experimental setup (as shown in Figure 4) and instructed to push directly into the force sensor i.e., along the ‘z’ axis, – either directly with the palm or via the gimbal device – using their right arm. They were tasked to generate their maximum isometric force for a total of 7 seconds. No visual feedback was provided about the amount or direction of the generated force. The beginning and end of each trial was communicated verbally by the experimenter.

The test was repeated in different conditions organized in the protocol presented in Figure 6. Each subject started by performing a single trial of maximum pushing force task (pre) using their preferred arm configuration. This was done to measure subjects’ initial performance. Then, each subject was randomly assigned to perform either the preferred – first – and the ‘undesired’ – after – arm configuration blocks, or vice versa. The preferred arm configuration allowed each subject to perform the pushing task in whatever posture they considered best for them. The ‘undesired’ arm configuration required subjects to push while maintaining what is generally considered an uncomfortable pushing configuration i.e., pushing with the wrist facing upward, maximizing pronation.

**Figure 6.**
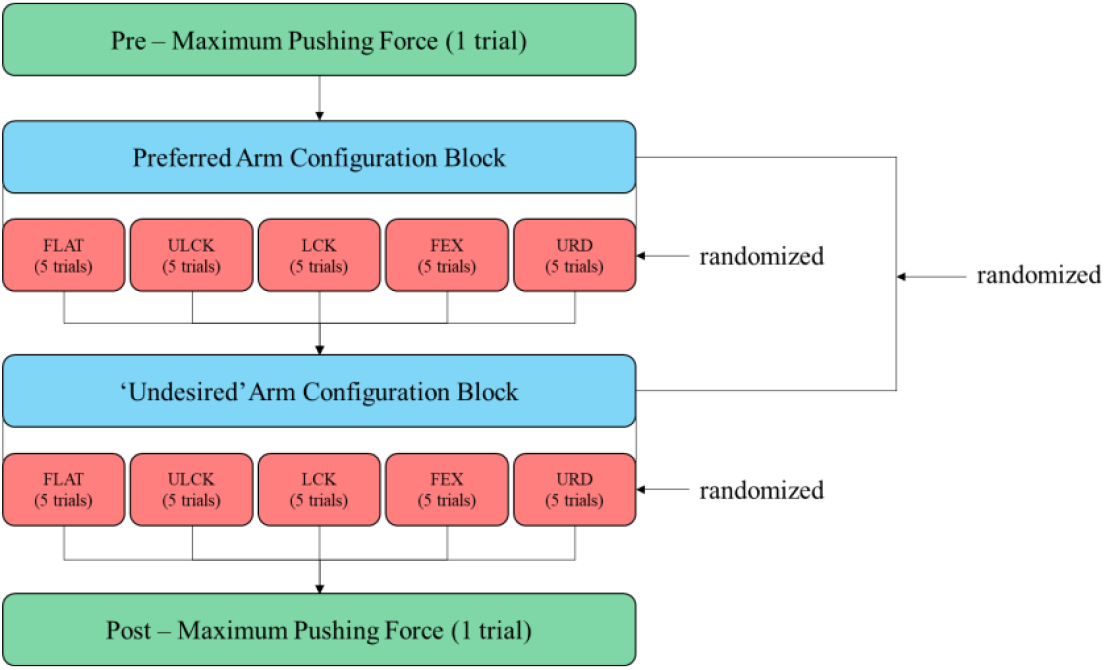
The experimental protocol. Each subject started by performing a single maximum pushing force trial (pre) in their preferred arm configuration. This was done to measure subjects’ initial performance. Then, each subject was randomly assigned to perform either the preferred. first. and the ‘undesired’. after. arm configuration blocks, or vice versa. The preferred arm configuration allowed each subject to perform the pushing task in whatever posture they considered best for them. The ‘undesired’ arm configuration required subjects to push while maintaining what is generally considered an uncomfortable pushing configuration i.e., pushing with the wrist facing upward, maximizing pronation. In each arm configuration block, subjects were tasked to perform the pushing task in 5 conditions: ‘FLAT’ i.e., pushing without the gimbal device with the palm of their hand; ‘ULCK’ i.e., pushing with the gimbal device while all axes were unlocked (a_1_, a_2_, a_3_); ‘LCK’ i.e., pushing with the gimbal device while all axes were locked; ‘FEX’ i.e., pushing with the gimbal device with flexion-extension axis (a_1_) unlocked and the others locked; ‘URD’ i.e., pushing with the gimbal device with both flexion-extension (a_1_) and ulnar-radial deviation (a^2^) axes unlocked. The 5 conditions were randomized between subjects and between blocks. Each condition was repeated 5 times. At the end of both arm configuration blocks, subjects performed a final (post) single trial maximum pushing force in their preferred arm configuration. This was done to account for the possible presence of fatigue or learning.

In each arm configuration block, subjects were tasked to perform the pushing task in 5 conditions: ‘FLAT’ i.e., pushing without the gimbal device using the palm of their hand; ‘ULCK’ i.e., pushing with the gimbal device while all axes were unlocked (*a*_1_, *a*_2_, *a*_3_); ‘LCK’ i.e., pushing with the gimbal device while all axes were locked; ‘FEX’ i.e., pushing with the gimbal device with only the flexion-extension axis (*a*_1_) unlocked and the others locked; ‘URD’ i.e., pushing with the gimbal device while both flexion-extension (*a*_1_) and ulnar-radial deviation (*a*^2^) axes were unlocked. The 5 conditions were randomized between subjects and between blocks. Each condition was repeated 5 times in succession (trials). Subjects were allowed to take breaks and rest at any time during the experiment.

At the end of both arm configuration blocks, subjects performed a final (post) single trial maximum pushing force task in their preferred arm configuration. This was done to account for the possible presence of fatigue or learning.

Overall, subjects performed a total of 50+2 pushing trials. The whole experiment took between 35 minutes to 1 hour – depending on subject breaks.

### Data Processing and Statistical Analysis

The generated force was computed as the magnitude of the force vector measured along the three translational axes of the force sensor:

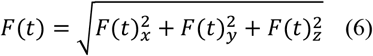

The force signal components (*F*(*t*)_*x*_, *F*(*t*)_*y*_, *F*(*t*)_*z*_) were sampled at 50 Hz using the ATI force/torque sensor. Out of the 7 seconds of signal recording, we considered a *T*_*w*_ = 3 *s* window, specifically, the portion of data going from 3 *s* ≤ *t* ≤ 6 *s*. This was the portion of signal clear of the initial force ramp-up as well as sufficiently away from the end to avoid any possible relaxation due to the expected termination of the trial^4^.

We decided to use as metric the magnitude of the force vector – rather than the ‘z’ direction force component – to have a measure of the total applied force independently of any eventual misalignment between the direction of force generation and the ‘z’ direction. A closer look at each force component revealed – as expected – that most of the force was concentrated along the ‘z’ direction, though some misalignment was observable. This further justified the decision to investigate the total force magnitude.

Two research questions were investigated in this work: (1) Is the maximum voluntary isometric force limited by the ability to generate stiffness during an unstable physical interaction task? (2) Does arm configuration influence maximum voluntary isometric force generation?

To statistically address these two research questions, we used a 2-way repeated-measures ANOVA design. We analyzed the effect of arm configuration (preferred, undesired), and pushing conditions (FLAT, ULCK, LCK, FEX, URD) on the generated steady-state maximum isometric force *F*(*t*). A significance threshold of *α* = 0.05 was used. Bonferroni corrected post-hoc t-tests were conducted in case of significant effects. The analysis was performed – on an individual subject basis – pooling each tested condition across the 5 trials. The decision to perform individual subjects’ analysis was taken to observe any individual preference in force generation. This was motivated by the Synergy Expansion hypothesis which indicates that different humans are not guaranteed to converge to the same motor coordination for the same motor task (Tessari et al., 2025).

An additional 2-way repeated-measures ANOVA was performed to assess the effect of individual subjects and time (pre- and post-) on the maximum voluntary isometric force. A significance threshold of *α* = 0.05 was considered. Bonferroni corrected post-hoc t-tests were conducted in case of significant effects. This additional test was included to investigate the presence of fatigue or learning in individual subjects.

As detailed in the Results section, since the differences due to arm configurations were quite small in amplitude, we also computed the median force difference ‘Δ*F*_*med*_’ (and relative error ‘Δ*e*_*med*_’) for each condition and trial, and then pooled them across subjects:

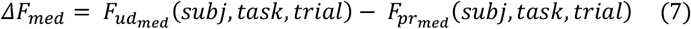

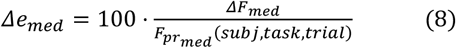

where 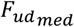 (*subj, task, trial*) represents the median force for a specific subject, condition, and trial in the undesired arm configuration, while 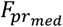 (*subj, task, trial*) represents the median force for a specific subject, condition, and trial in the preferred arm configuration.

The distribution of median force differences and median relative errors across subjects was compared with humans’ ‘just noticeable difference’ (JND) threshold. JND is the measure of the minimum difference between two stimuli that is necessary in order for the difference to be reliably perceived. According to Allin et al., JND is relatively constant over a range of different applied forces with values between 2.5 N and 10 N (Allin et al., 2002). This further analysis was performed to assess whether the observed differences between arm configurations went beyond what humans could actually perceive.

## Results

### Representative Subject Force Trajectory

Figure 7 shows the total force magnitude as well as the 3 different force sub-components (*F*_*x*_, *F*_*y*_, *F*_*z*_) for a representative subject across all trials and conditions. All subjects were able to complete the experiment. All but one subject (Subj. 9) exhibited a very similar behavior to the one observed in Figure 7. Subject 9 also did not complete the ‘LCK’ condition in the ‘undesired’ arm configuration due to a personal time constraint. Detailed results for all subjects are reported in the Supplementary Materials.

**Figure 7.**
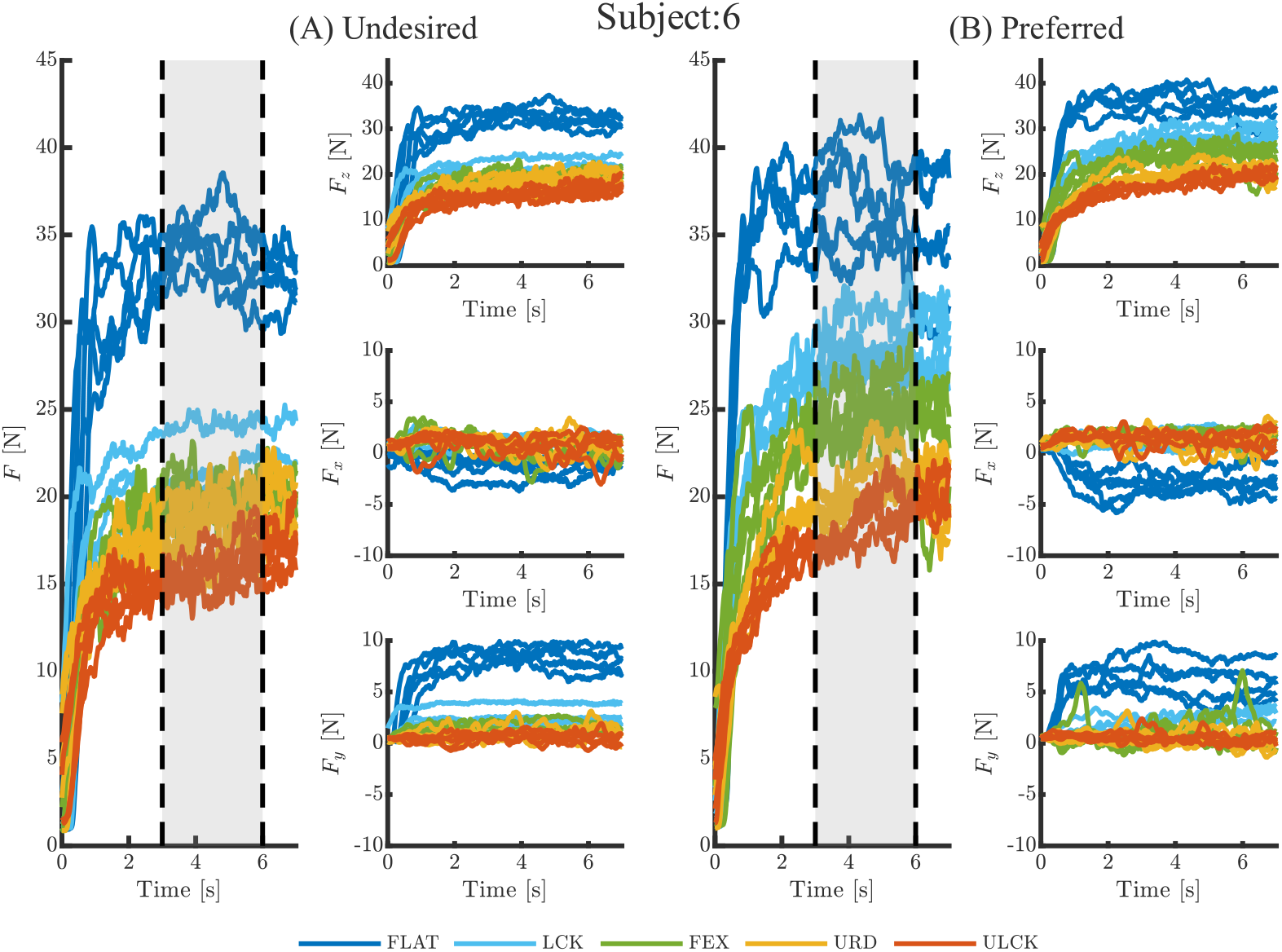
Force temporal trajectories for a representative subject (nr. 6). Panel (A) shows the total force magnitude ‘F’ (left column), and each of the three sub-components: ‘F_z_’ top right, ‘F_x_’ middle right, ‘F_y_’ bottom right columns, for the undesired hand configuration. Panel (B) shows the total force magnitude ‘F’ (left column), and each of the three sub-components: ‘F_z_’ top right, ‘F_x_’ middle right, ‘F_y_’ bottom right columns, for the preferred hand configuration. In each plot the different pushing conditions are presented in different solid color lines: FLAT in dark blue, LCK in light blue, FEX in green, URD in yellow, and ULCK in red. Each individual line represents a single trial. In the total force magnitude plots, two dashed-black lines delimit a shaded grey area representing the time window considered for the analysis of maximum voluntary force performance.

It is observable (from Figure 7) that the total force magnitude ‘*F*’ ramped up from zero to near a steady-state value at about 3 seconds, and was maintained for the rest of the task. It is also notable that the total force magnitude and the force along the ‘z’ axis were very similar both in terms of temporal evolution as well as magnitude. The forces along the other two orthogonal axes (‘x’ and ‘y’) are considerably smaller. Moreover, these forces oscillated around zero for all conditions except the FLAT one. This is expected, since in all cases – except the FLAT condition – a ball joint connected the gimbal with the sensor, which made the generation of lateral forces much harder. More importantly, a pronounced decrease in total force is observable when moving across conditions from FLAT to ULCK.

### The Effect of Pushing Condition and Arm Configuration on Maximum Force

Subjects presented a diverse range of forces across conditions. The strongest subject was number 5 with a maximum median force of 36.5 N, while the weakest subject was number 9 with a maximum median force of 13.4 N – almost a 3:1 ratio in performance. Despite the diversity in force generation ability, all subjects experienced across conditions an overall loss of performance in terms of maximum voluntary isometric force generation – independently of arm configuration. Detailed results are reported in Table 1. On average, subjects lost 37.7 ± 9.2% of maximum voluntary isometric force going from the most stable (FLAT) to the most unstable (ULCK – fully unlocked) configurations. Subject 6 presented the most pronounced performance loss (53.7%) in the undesired arm configuration; while subject 9 presented the least performance loss (23.1%) also in the undesired arm configuration.

**Table 1.** Median forces generated in each task condition (FLAT, LCK, FEX, URD, ULCK), arm configuration (Undesired, Preferred), for every subjects. The last two columns also report the maximum difference between median forces between task conditions, as well as the maximum percentage difference between median forces normalized to the maximum median force measure in each subject and arm configuration. The maximum and minimum percentage differences among subjects are bolded and colored in red and blue, respectively.

| Subj. | Arm | $F_{med_{FLAT}}$<br>[N] | $F_{med_{LCK}}$<br>[N] | $F_{med_{FEX}}$<br>[N] | $F_{med_{URD}}$<br>[N] | $F_{med_{ULCK}}$<br>[N] | $\Delta F_{med_{max}}$<br>[N] | Perc.<br>Diff.<br>[%] |
| --- | --- | --- | --- | --- | --- | --- | --- | --- |
| 1 | Undesired | 24.49 | 19.27 | 15.53 | 14.45 | 14.05 | 10.44 | 42.64 |
| 1 | Preferred | 26.04 | 20.79 | 17.47 | 15.75 | 12.95 | 13.09 | 50.27 |
| 2 | Undesired | 30.76 | 25.09 | 19.46 | 20.26 | 17.14 | 13.62 | 44.28 |
| 2 | Preferred | 31.23 | 23.62 | 21.17 | 20.18 | 19.94 | 11.29 | 36.15 |
| 3 | Undesired | 18.10 | 16.01 | 13.94 | 12.88 | 13.57 | 5.22 | 28.86 |
| 3 | Preferred | 19.95 | 17.39 | 17.03 | 15.14 | 15.90 | 4.81 | 24.12 |
| 4 | Undesired | 26.74 | 21.97 | 20.43 | 19.38 | 19.01 | 7.72 | 28.89 |
| 4 | Preferred | 27.01 | 20.89 | 19.74 | 19.17 | 18.68 | 8.33 | 30.85 |
| 5 | Undesired | 33.80 | 26.79 | 25.44 | 20.31 | 20.79 | 13.49 | 39.91 |
| 5 | Preferred | 36.49 | 25.46 | 22.89 | 20.81 | 21.32 | 15.69 | 42.98 |
| 6 | Undesired | 33.93 | 20.67 | 19.71 | 18.65 | 15.72 | 18.21 | **53.68** |
| 6 | Preferred | 36.40 | 27.93 | 24.54 | 20.17 | 18.15 | 18.25 | 50.14 |
| 7 | Undesired | 20.91 | 14.43 | 12.29 | 12.08 | 11.36 | 9.55 | 45.67 |
| 7 | Preferred | 18.99 | 15.22 | 13.92 | 11.37 | 10.87 | 8.11 | 42.73 |
| 8 | Undesired | 30.56 | 27.46 | 20.51 | 19.35 | 22.05 | 11.21 | 36.69 |
| 8 | Preferred | 33.12 | 28.22 | 24.78 | 23.11 | 24.09 | 10.01 | 30.23 |
| 9 | Undesired | 11.61 | N.A. | 13.40 | 11.29 | 10.31 | 3.10 | **23.10** |
| 9 | Preferred | 13.79 | 17.18 | 16.11 | 13.05 | 12.56 | 4.62 | 26.90 |
| **Avg.** |  |  |  |  |  |  |  | **37.67** |

Figure 8 presents boxplots of the total force ‘*F*’ during the selected time window (3 *s* ≤ *t* ≤ 6 *s*) for each subject, task condition, and arm configuration. The 2-way repeated measure ANOVA analysis found a significant effect of task condition (FLAT, LCK, FEX, URD, ULCK), a significant effect of arm configuration (undesired, preferred), and a significant interaction for all subjects. Bonferroni corrected post-hoc t-tests revealed that 94.7% of all the pair-wise comparisons were significant. The non-significant pair-wise comparisons – reported graphically in Figure 8 –predominantly (∼60%) involved arm configuration pair-wise comparisons. The complete details of the statistical analysis are reported in the Supplementary Materials.

**Figure 8.**
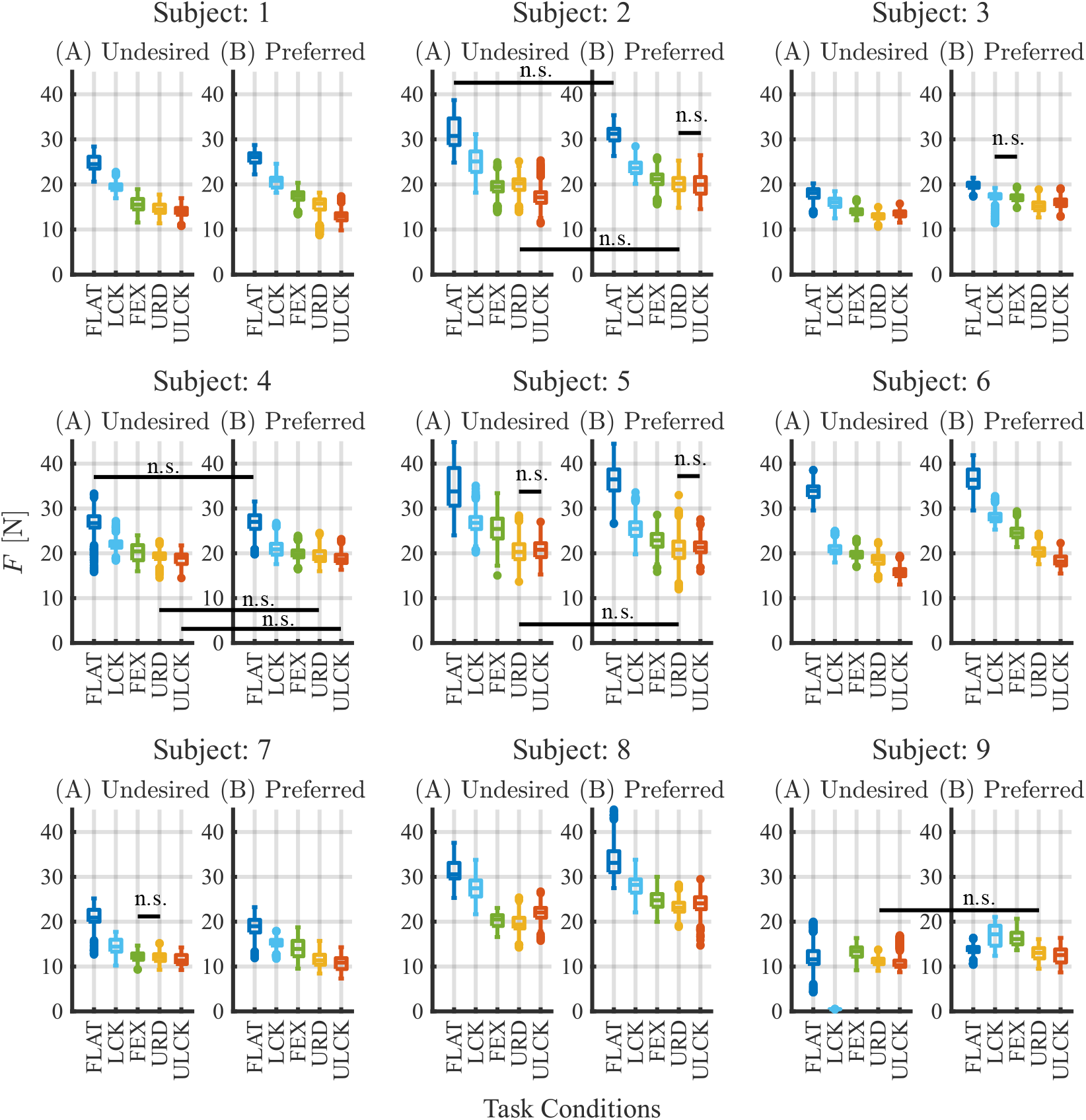
Collection of subjects individual boxplots showing the distribution of generated forces in each task condition (FLAT, LCK, FEX, URD, ULCK) and arm configuration: (A) Undesired, (B) Preferred. Non-significant differences are also reported using horizontal black lines. The task conditions are reported using the same color pattern throughout the paper.

Each condition, moving from FLAT to ULCK, progressively complicated the stabilization of the task by reducing the wrist’s ability to generate torque in specific directions. When going from FLAT to LCK, the task goes from being a stable pushing task to an unstable one i.e., pushing on a stick. Then, when moving from LCK to FEX, subjects lose their ability to generate wrist flexion-extension torque. In the URD case, subjects additionally lose their ability to generate wrist ulnar-radial deviation torque. At last, in the ULCK case, subjects lose their ability to generate wrist torque also in pronation-supination.

It is interesting to observe how the maximum voluntary force has a significantly and almost-monotonically decreasing trend in all subjects and in both arm configurations when going from the FLAT condition i.e., pushing directly against the robot, to the ULCK one i.e., pushing in the most challenging condition with all the gimbal degrees of freedom unlocked. However, exceptions in some specific cases are observable. For example, subject 9 has a non-monotonic behavior between the FLAT and LCK conditions. However, the force still shows a significantly decreasing maximum force going from the fully-locked (LCK) to the fully-unlocked (ULCK) gimbal conditions.

Our results demonstrate that going from a stable to an unstable pushing task (FLAT **→** LCK) significantly reduced the maximum voluntary isometric force performance. By continuously impairing the wrist’s ability to stabilize the task (LCK **→** FEX **→** URD **→** ULCK), subjects continued to significantly reduce their maximum voluntary isometric force. The effect of removing the wrist’s ability to generate torque decreased non-linearly with an approximately exponentially decaying trend i.e., the more wrist ability to generate torque was removed the less the maximum voluntary isometric force was reduced. This was evident in all subjects and clearly observable in Figure 8. We refrained from interpolating the data between task-conditions since the scale is ordinal but the intervals’ distance cannot be numerically defined.

As mentioned previously, arm configuration was found to have a significant effect on maximum force performance. However, as observable in Figure 8, the use of the two different arm configurations did not seem to substantially change the amount of generated force in each condition. To better highlight this, we computed the median force difference Δ*F*_*med*_ and the relative percentage error Δ*e*_*med*_ between arm configurations in each task, condition, and subject. (Please refer to the Materials and Methods section for more details on the calculation of these two metrics). Figure 9 presents the histograms of Δ*F*_*med*_ and Δ*e*_*med*_.

**Figure 9.**
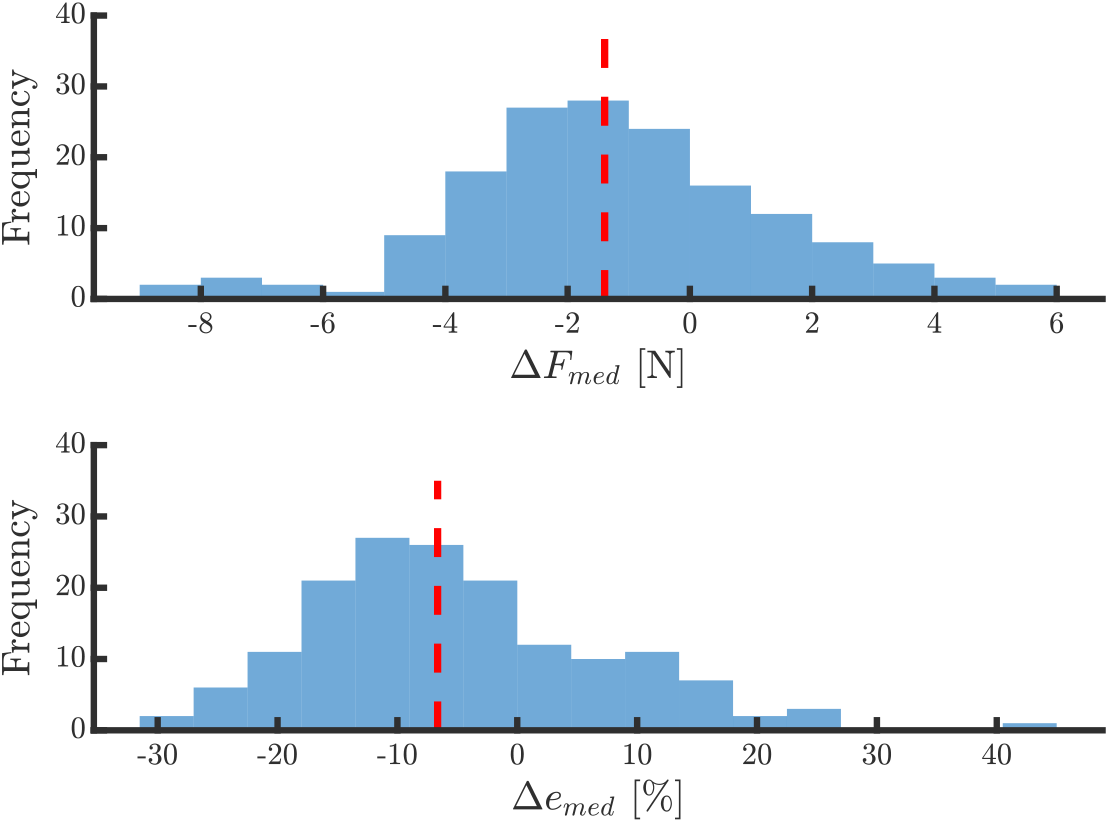
Histograms of the median differences in arm configuration between task conditions pooled across subjects. The top plot shows the histogram for the median force difference ΔF_med_. The bottom plot presents the relative percentage error Δe_med_ computed with respect to the preferred arm configuration. Please refer to the Material & Methods section for details. The vertical dashed red lines represent the median of the distributions: median(ΔF_med_) = −1.39 N, median(Δe_med_) = −6.64%.

It can be seen that the distribution of force differences between the undesired and preferred arm configuration is unimodal and almost centered around zero (*median*(Δ*F*_*med*_) = −1.39 *N, median*(Δ*e*_*med*_) = −6.64%). Subjects were – on average – pushing slightly harder in the preferred arm configuration, with almost symmetric results with respect to zero. Moreover, it is worth emphasizing that the median difference in forces between arm configurations was well below the Just Noticeable Difference range, reported to be 2.5 to 10 N (Allin et al., 2002). This suggests that – even if the difference in maximum voluntary isometric force between arm configurations was significant – its effect was probably negligible since it was below the range of perception of differences typically observed in humans. In other words, the effect of arm configuration was negligible compared to the effect of task condition.

### Pre-post Maximum Voluntary Force

Figure 10 reports the boxplots of the maximum voluntary isometric force measured pre- and post-the main experiment. A two-way repeated measure ANOVA found a significant effect of subject, a significant effect of time (pre vs. post), and a significant interaction. Bonferroni corrected post-hoc pairwise comparisons demonstrated a significant difference between pre- and post-measurements in all subjects except subject 4.

**Figure 10.**
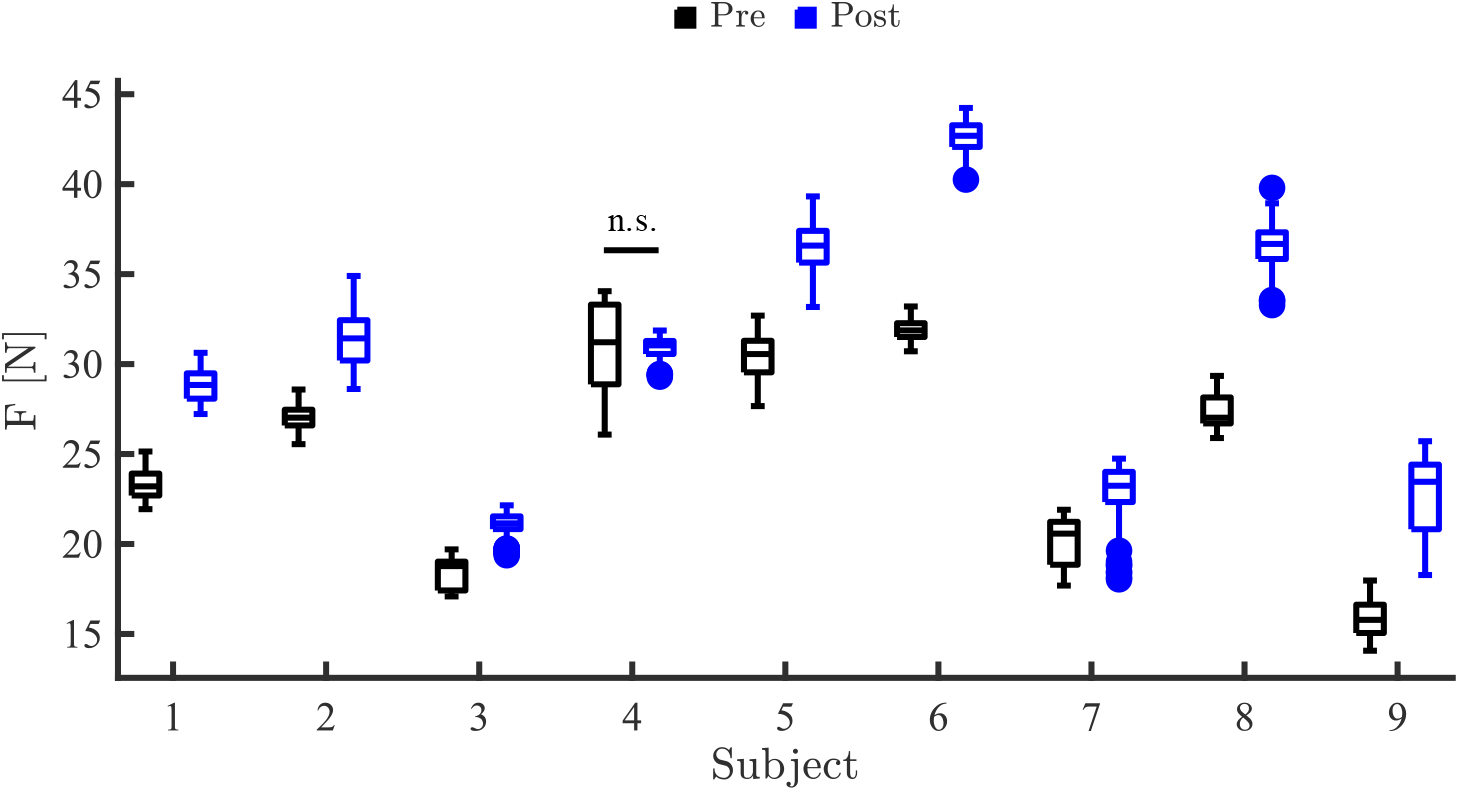
Boxplot presenting a comparison between the maximum voluntary isometric force measured pre- and post-the experiment. The black boxplots represent the pre-test, while the blue boxplots show the post-test. A horizontal solid line is added to show non-significant pair-wise comparison.

Excluding subject 4, whose performance remained constant, all the other subjects substantially improved their maximum voluntary isometric force i.e., their pushing ability. The best improvement was obtained by subject 9 who increased the maximum pushing force by +48.6%, while the smallest improvement was achieved by subject 3 (+12.7%). On average, subjects (excluding subj. 4) increased their pushing performance by +25.5%.

This was an unexpected result, as it suggests that: (i) subjects did not seem to show any measurable effect of fatigue, and (ii) subjects learned to push harder in less than an hour of experimental experience.

## Discussion

In this paper we investigated two research questions: (i) is human maximum force production limited by the ability to generate stabilizing stiffness? (ii) does arm configuration influence the maximum force generation?

To answer these two research questions, we developed an experiment in which subjects were asked to perform a simple physical interaction task i.e., pushing as hard as possible. The pushing task was performed in both stable (FLAT) and unstable conditions (LCK – fully locked, FEX – flexion/extension unlocked, URD – ulnar/radial deviation and flexion/extension unlocked, ULCK – fully unlocked). Due to the use of an unlockable gimbal apparatus, the unstable conditions progressively decreased the ability to generate a stabilizing action at the wrist. These conditions were tested in both the subjects’ preferred arm configuration, as well as in a highly undesirable arm configuration i.e., pushing while maintaining the wrist facing upward (maximum pronation). This experiment drew inspiration from the work by (Rancourt & Hogan, 2009) in which they showed preliminary evidence that force production during physical interaction is affected by the type of tool, arm configuration, and performed task. Their results motivated our in-depth exploration of the effect of stabilization and individual preferences on force production.

Our analysis showed that the maximum pushing force significantly decreased when moving from a stable to an unstable pushing condition. The arm configuration remained essentially constant. Consequently, the moment arms relating muscle tension to externally exerted force were unchanged. Yet the maximum force decreased with an approximately exponential trend (Figure 8) as the unusable wrist degrees of freedom (DoF) increased.

Studying the effect of extreme changes of arm configuration, we did find a significant effect of arm configuration in all subjects and most conditions. However, the magnitude of this effect was well below the threshold of Just Noticeable Difference of human perception. The negligible effect of extreme variations of pose confirms that the skeletal configuration had a minimal role in the observed drop in maximum pushing performance, which has to be attributed to a loss in ability to stabilize the task by generating stiffness.

We know from biomechanics that unstable pushing tasks require a stabilizing stiffness to generate force (see Materials & Methods, as well as (Rancourt & Hogan, 2009)). By removing directions in which the wrist could generate stabilizing torque, we experimentally observed a decrease in pushing performance. This indicates that stiffness is the limiting factor in the generation of force i.e., if we can’t generate enough stiffness, we will not be able to generate enough output force. Interestingly, in the pushing task considered, the wrist, and the muscles actuating it, cannot significantly contribute to the force generation task, especially if the hand and forearm are collinear and aligned with the main pushing direction. They only contribute to stabilization. Moreover, if the wrist was a perfect ball-joint or U-joint – though we know that it isn’t (Charles & Hogan, 2011) – then we would expect that disabling one wrist DoF would eliminate all wrist muscle contributions to stiffness. However, most subjects presented significant differences in maximum pushing force by incrementally disabling the wrist’s ability to generate torque, thus further confirming a significant role of each wrist DoF to stabilization and, consequently, force generation.

In principle, a loss of stabilization at the wrist can be compensated by other joints e.g., the shoulder, but this compensation is evidently not enough. Muscles that were previously used for pushing, may now need to be used for stabilizing the task. The overall result is a drop in performance i.e., the maximum output force decreases. On a separate – but relevant – note, it is worth underlining that individual skeletal muscle activity tends to de-stabilize our endo-skeleton, even with no external force production. This is a direct consequence of biomechanics and the fact that muscles can only exert tensile forces and not compressive ones.

The human musculoskeletal system – thanks to its redundancy – offers multiple solutions to the same motor task, with negligible or less-than-noticeable differences (Tessari et al., 2025). This applies also to our very simple pushing task, in which the preferred and undesired arm configurations led to comparable performance.

Our results do not mean that subjects will not have a preferred way of pushing. In fact, most subjects’ preferred arm configuration for pushing was with the wrist facing downwards. This might be due to other musculoskeletal constraints or preferences not investigated in this study. However, even when looking at subjects’ preferred limb configuration and posture for pushing, we could clearly see differences between the analyzed subjects. These differences were not quantified but they further emphasize the versatility of the human musculoskeletal system.

We also investigated the effect of time (pre-vs. post-) on the maximum voluntary isometric force exertion. This was initially introduced to test the possible functional effects of fatigue. Surprisingly, the pre-post analysis showed that all subjects but one significantly *improved* their pushing performance (on average by +25.5%) at the end of the experiment. This was despite the fact that most subjects self-reported some level of muscle fatigue. How can we explain this? One hypothesis is that during the pushing task two opposing processes were occurring. A fatiguing process, which should reduce overall performance, and a learning process which might improve overall performance. In our experiment, the difference between the processes was likely dominated by the learning.

Future studies should explore the interesting trade-off between these two opposing processes in both simple and complex physical interaction tasks. Moreover, it might also be interesting to further explore humans’ preference to perform tasks in what we here define as a “motor fovea” i.e., the human preference to perform upper limb tasks half meter below the shoulder and about a half meter in front of the sternum. Manipulability measures from the world of robotics – such as the one developed by Lachner et al. (Lachner et al., 2020) or Yoshikawa (Yoshikawa, 1985b, 1985a) – could be adapted to study human preferences in the selection of skeletal configurations for motion and physical interaction.

Our study may inform therapeutic treatments aimed at restoring motor functions. For example, if generating stabilizing stiffness is a higher priority, maybe we should focus on restoring this ability before teaching users how to generate force. Additionally, our findings may also affect the way we design control strategies for assistive and rehabilitative robots e.g., we should focus on control strategies that consider stabilization rather than just force generation.

This research highlights the important fact that muscles are not just force-producers. External factors determine whether muscle activation yields force or motion or something in-between. In contrast, muscle activation always increases stiffness or, more generally, mechanical impedance.

In conclusion, our results confirm that in some important functional contexts force generation is limited by our ability to generate stiffness. This holds true across subjects, even if multiple – equally feasible – preferences in skeletal configuration were used to perform the same motor task i.e., pushing.

## Supplementary Materials

### Subjects’ Force Trajectories

The total force ‘*F*’ and force sub-components (*F*_*x*_, *F*_*y*_, *F*_*z*_) temporal trajectories during the experiment for all the subjects (1 to 9) are reported in the following supplementary figures.

**Figure 11.**
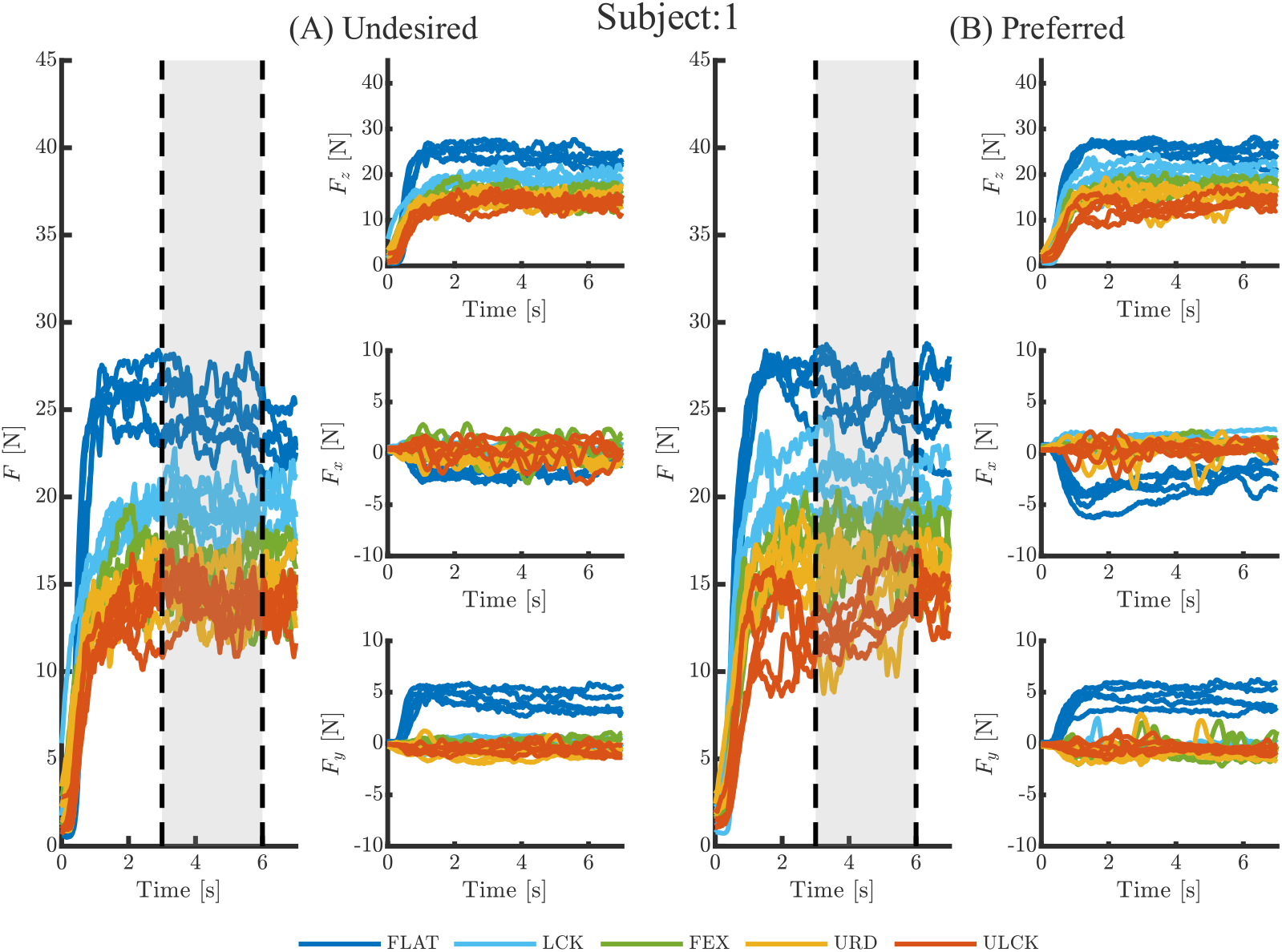
Force temporal trajectories for Subject 1. Panel (A) shows the total force magnitude ‘F’ (left column), and each of the three sub-components: ‘F_z_’ top right, ‘F_x_’ middle right, ‘F_y_’ bottom right columns, for the undesired arm configuration. Panel (B) shows the total force magnitude ‘F’ (left column), and each of the three sub-components: ‘F_z_’ top right, ‘F_x_’ middle right, ‘F_y_’ bottom right columns, for the preferred arm configuration. In each plot the different pushing conditions are presented in different solid color lines: FLAT in dark blue, LCK in light blue, FEX in green, URD in yellow, and ULCK in red. Each individual line represents a single trial. In the total force magnitude plots, two dashed-black lines delimit a shaded grey area representing the considered time window for the analysis of maximum voluntary force performance.

**Figure 12.**
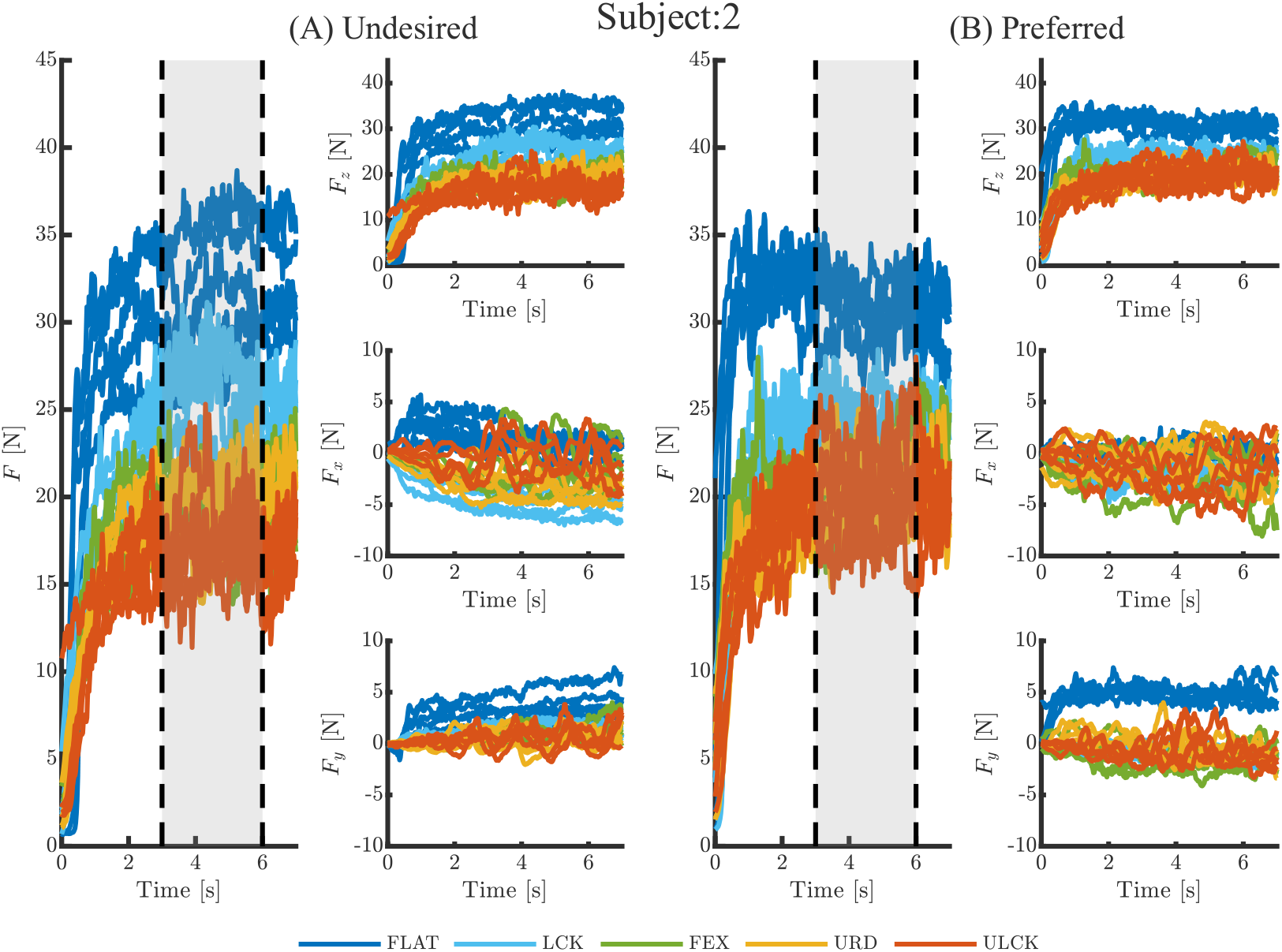
Force temporal trajectories for Subject 2. Panel (A) shows the total force magnitude ‘F’ (left column), and each of the three sub-components: ‘F_z_’ top right, ‘F_x_’ middle right, ‘F_y_’ bottom right columns, for the undesired arm configuration. Panel (B) shows the total force magnitude ‘F’ (left column), and each of the three sub-components: ‘F_z_’ top right, ‘F_x_’ middle right, ‘F_y_’ bottom right columns, for the preferred arm configuration. In each plot the different pushing conditions are presented in different solid color lines: FLAT in dark blue, LCK in light blue, FEX in green, URD in yellow, and ULCK in red. Each individual line represents a single trial. In the total force magnitude plots, two dashed-black lines delimit a shaded grey area representing the considered time window for the analysis of maximum voluntary force performance.

**Figure 13.**
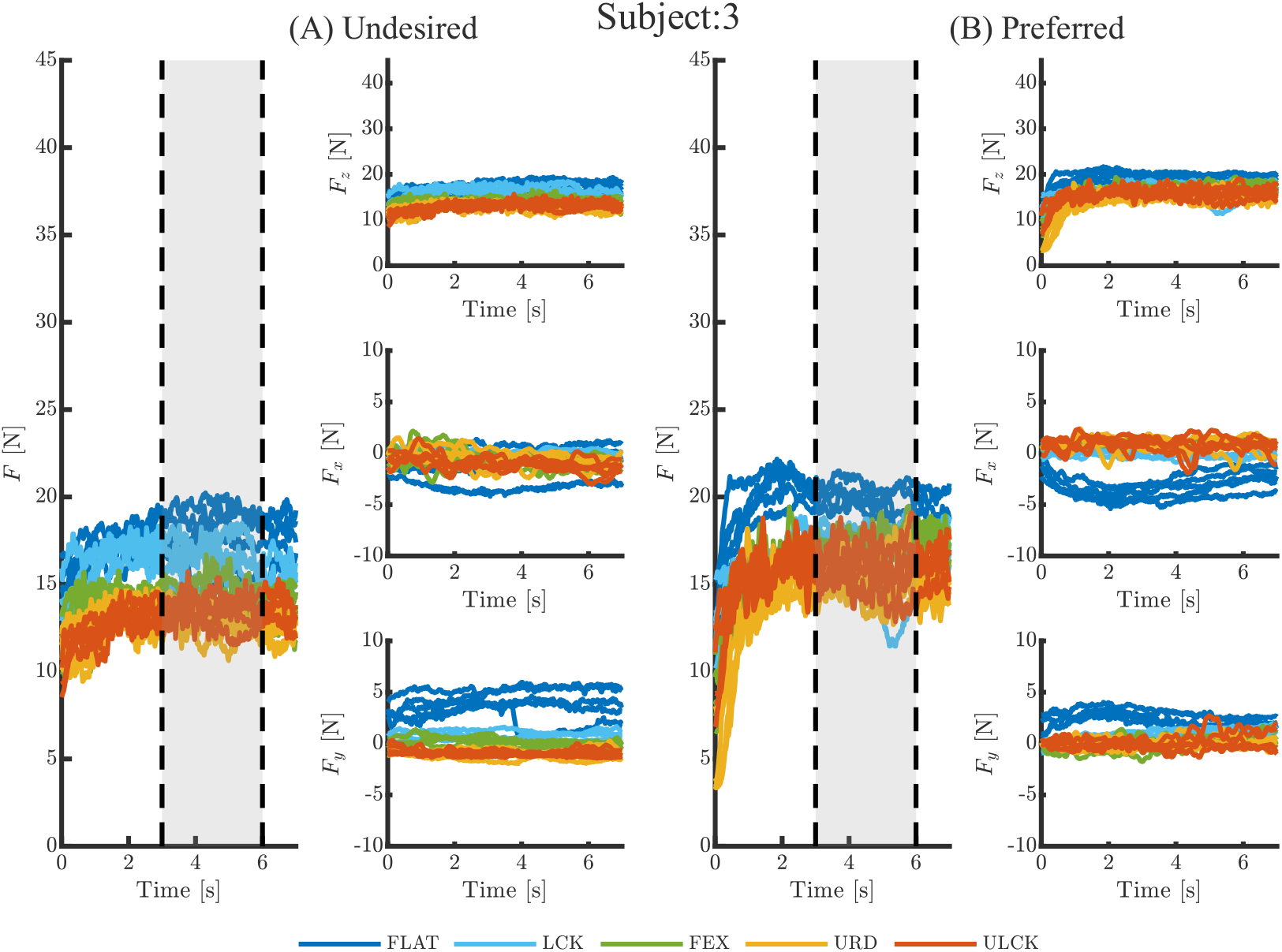
Force temporal trajectories for Subject 3. Panel (A) shows the total force magnitude ‘F’ (left column), and each of the three sub-components: ‘F_z_’ top right, ‘F_x_’ middle right, ‘F_y_’ bottom right columns, for the undesired arm configuration. Panel (B) shows the total force magnitude ‘F’ (left column), and each of the three sub-components: ‘F_z_’ top right, ‘F_x_’ middle right, ‘F_y_’ bottom right columns, for the preferred arm configuration. In each plot the different pushing conditions are presented in different solid color lines: FLAT in dark blue, LCK in light blue, FEX in green, URD in yellow, and ULCK in red. Each individual line represents a single trial. In the total force magnitude plots, two dashed-black lines delimit a shaded grey area representing the considered time window for the analysis of maximum voluntary force performance.

**Figure 14.**
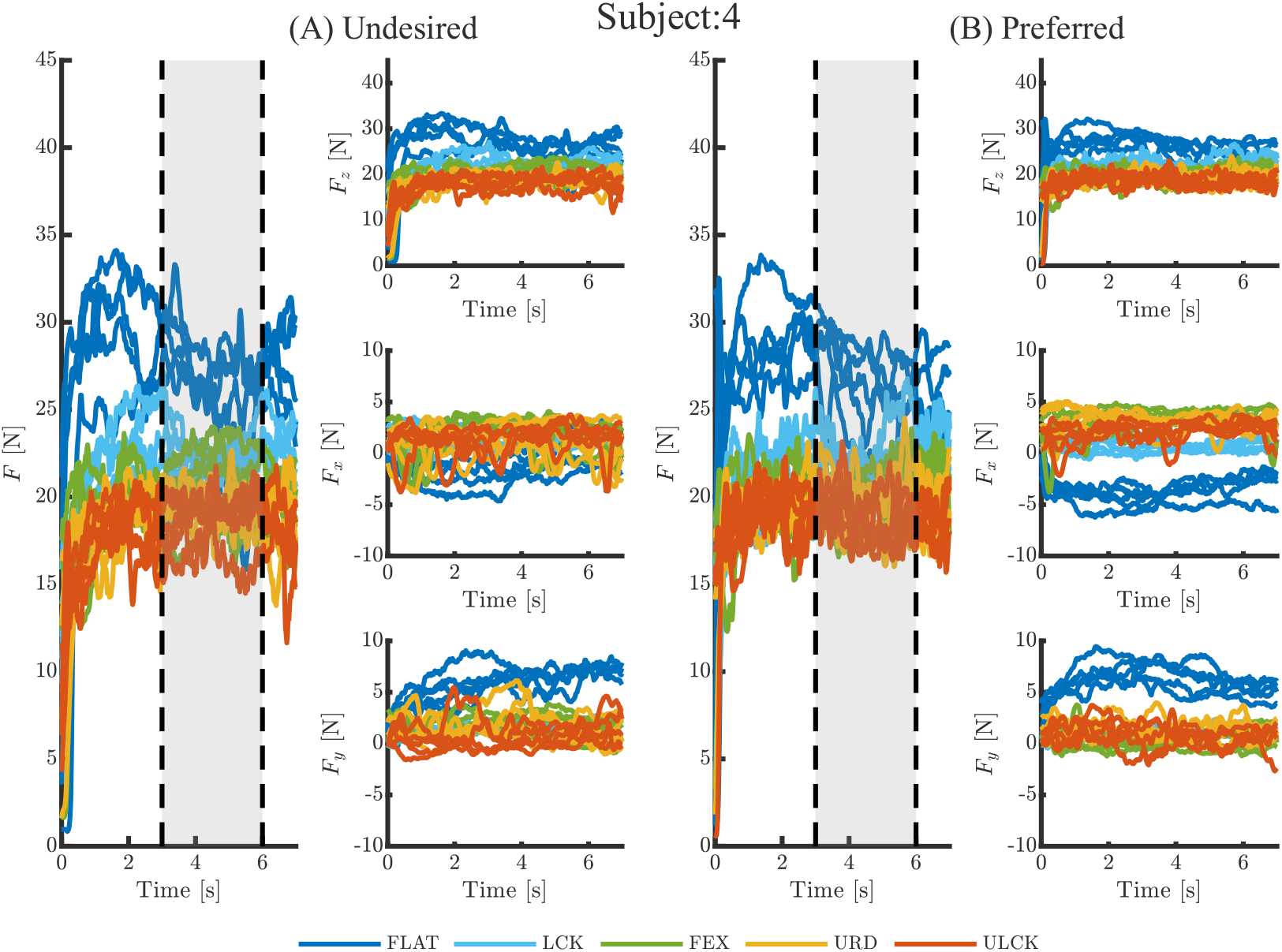
Force temporal trajectories for Subject 4. Panel (A) shows the total force magnitude ‘F’ (left column), and each of the three sub-components: ‘F_z_’ top right, ‘F_x_’ middle right, ‘F_y_’ bottom right columns, for the undesired arm configuration. Panel (B) shows the total force magnitude ‘F’ (left column), and each of the three sub-components: ‘F_z_’ top right, ‘F_x_’ middle right, ‘F_y_’ bottom right columns, for the preferred arm configuration. In each plot the different pushing conditions are presented in different solid color lines: FLAT in dark blue, LCK in light blue, FEX in green, URD in yellow, and ULCK in red. Each individual line represents a single trial. In the total force magnitude plots, two dashed-black lines delimit a shaded grey area representing the considered time window for the analysis of maximum voluntary force performance.

**Figure 15.**
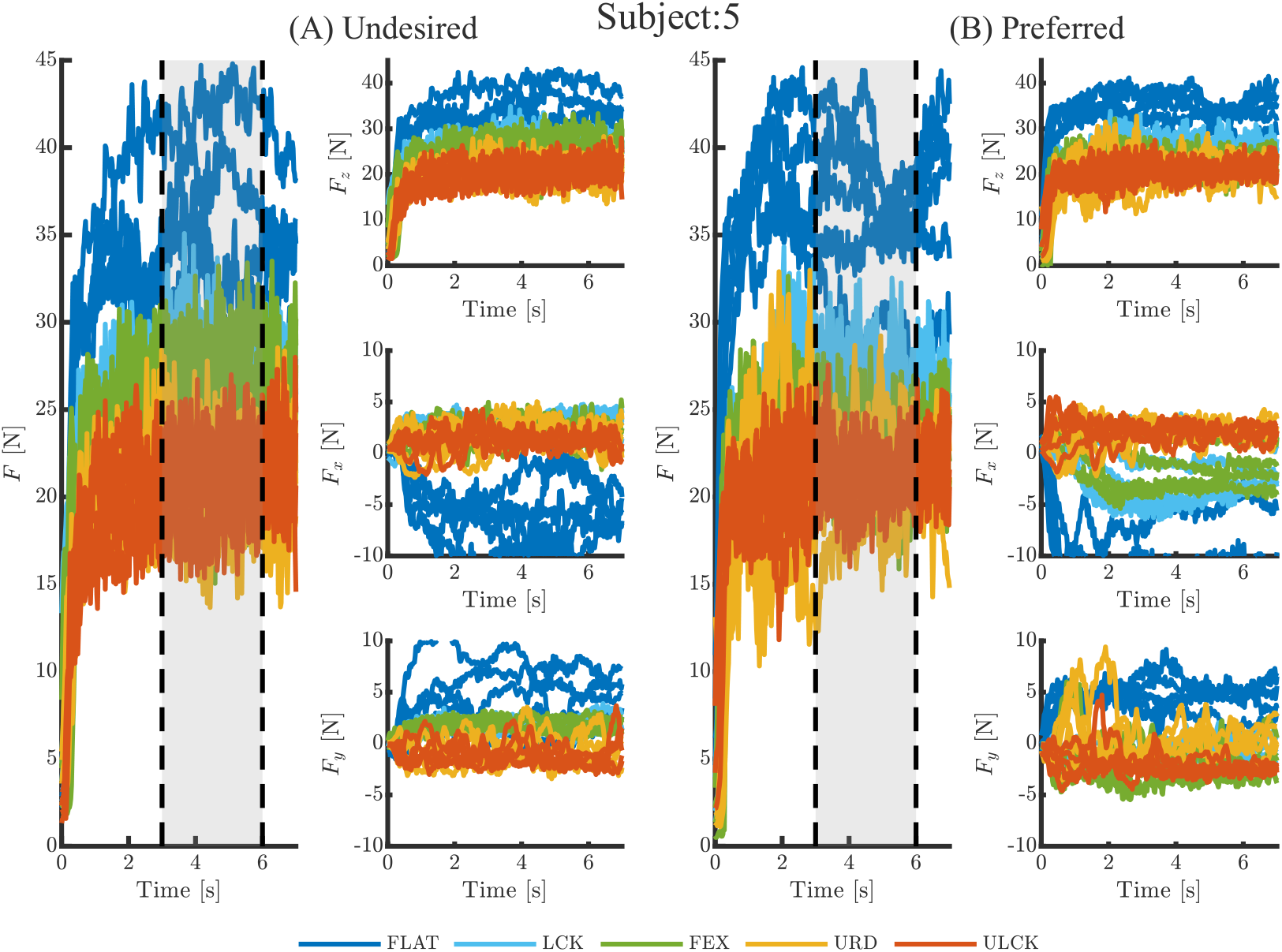
Force temporal trajectories for Subject 5. Panel (A) shows the total force magnitude ‘F’ (left column), and each of the three sub-components: ‘F_z_’ top right, ‘F_x_’ middle right, ‘F_y_’ bottom right columns, for the undesired arm configuration. Panel (B) shows the total force magnitude ‘F’ (left column), and each of the three sub-components: ‘F_z_’ top right, ‘F_x_’ middle right, ‘F_y_’ bottom right columns, for the preferred arm configuration. In each plot the different pushing conditions are presented in different solid color lines: FLAT in dark blue, LCK in light blue, FEX in green, URD in yellow, and ULCK in red. Each individual line represents a single trial. In the total force magnitude plots, two dashed-black lines delimit a shaded grey area representing the considered time window for the analysis of maximum voluntary force performance.

**Figure 16.**
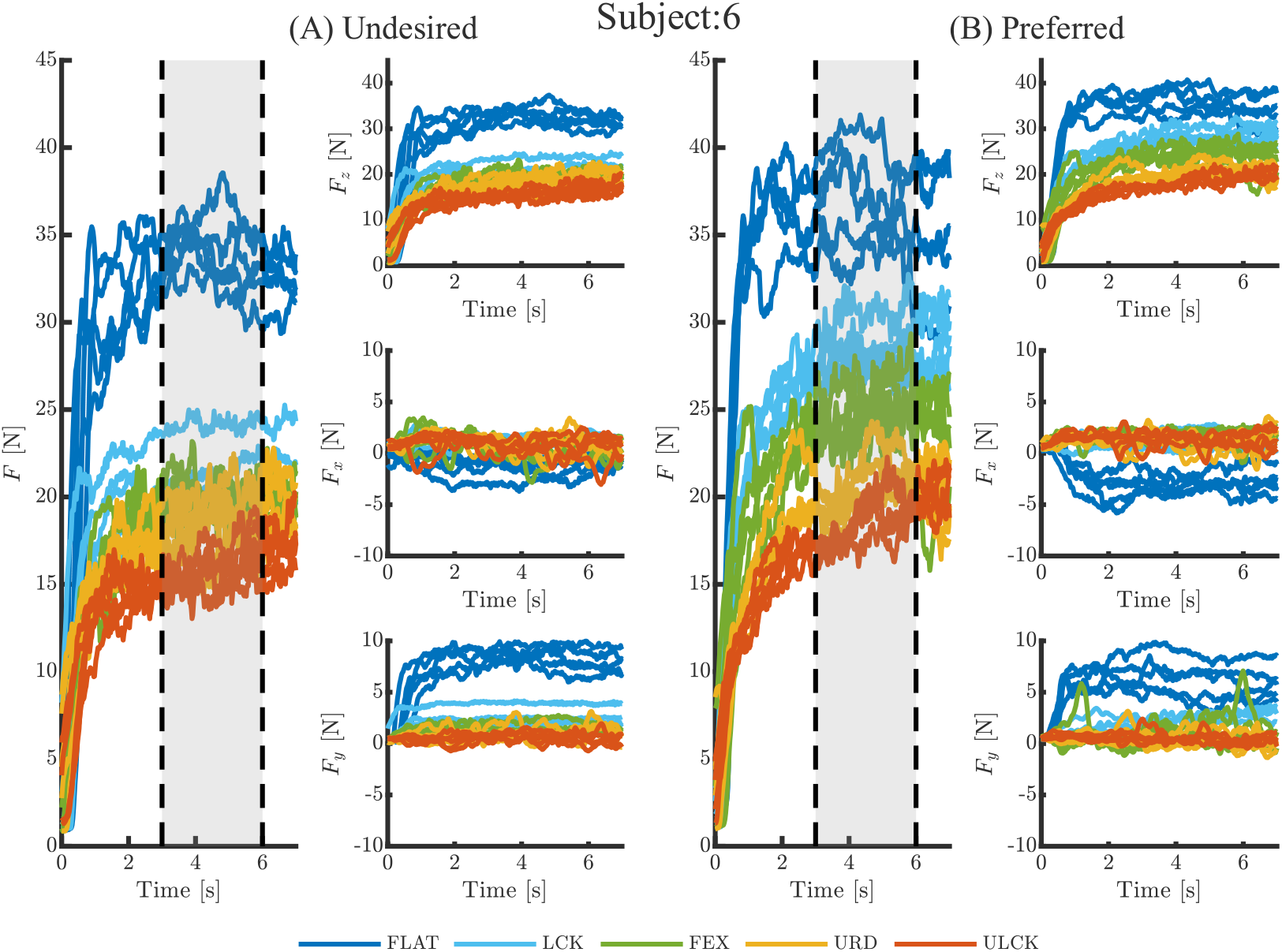
Force temporal trajectories for Subject 6. Panel (A) shows the total force magnitude ‘F’ (left column), and each of the three sub-components: ‘F_z_’ top right, ‘F_x_’ middle right, ‘F_y_’ bottom right columns, for the undesired arm configuration. Panel (B) shows the total force magnitude ‘F’ (left column), and each of the three sub-components: ‘F_z_’ top right, ‘F_x_’ middle right, ‘F_y_’ bottom right columns, for the preferred arm configuration. In each plot the different pushing conditions are presented in different solid color lines: FLAT in dark blue, LCK in light blue, FEX in green, URD in yellow, and ULCK in red. Each individual line represents a single trial. In the total force magnitude plots, two dashed-black lines delimit a shaded grey area representing the considered time window for the analysis of maximum voluntary force performance.

**Figure 17.**
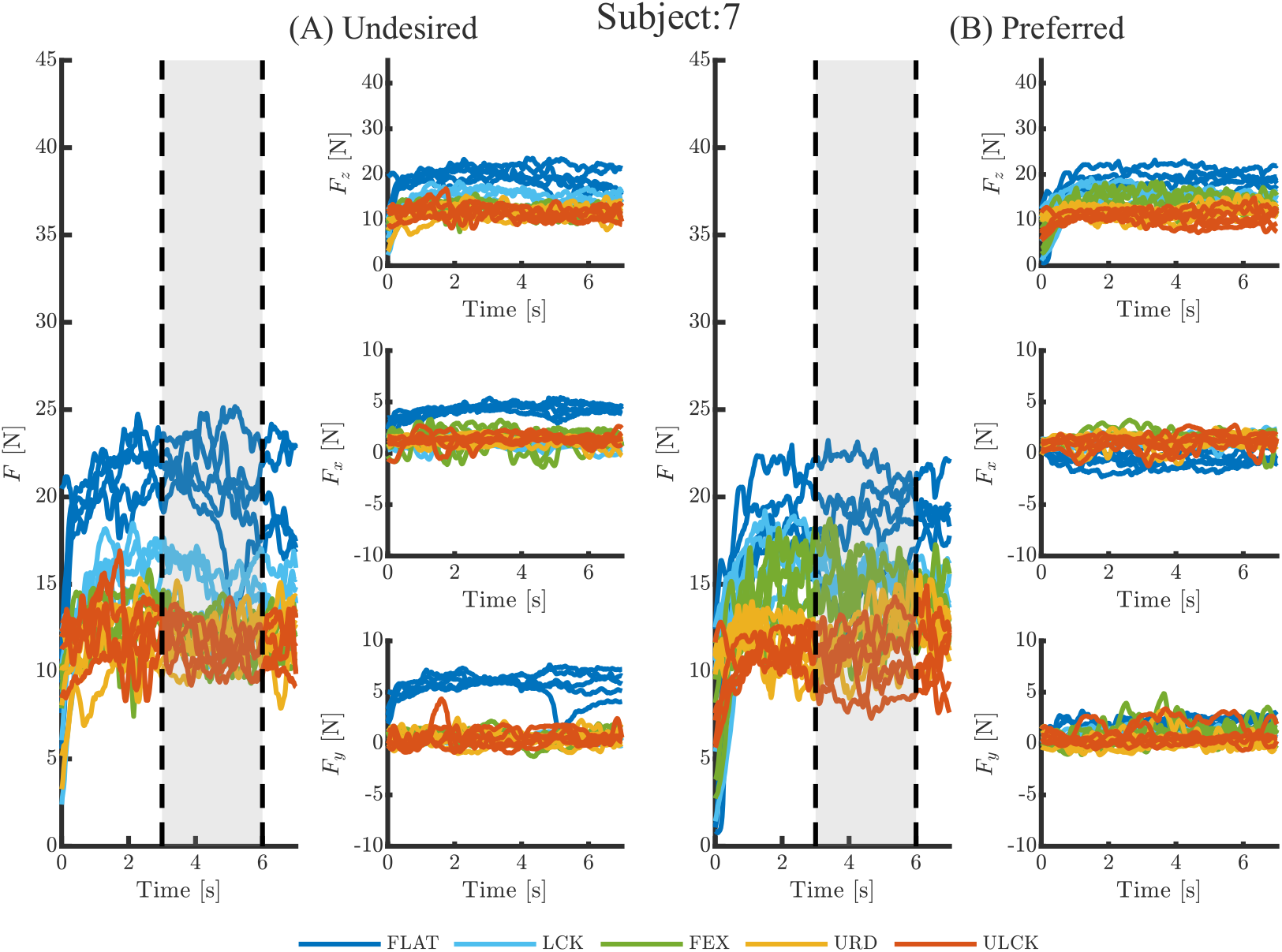
Force temporal trajectories for Subject 7. Panel (A) shows the total force magnitude ‘F’ (left column), and each of the three sub-components: ‘F_z_’ top right, ‘F_x_’ middle right, ‘F_y_’ bottom right columns, for the undesired arm configuration. Panel (B) shows the total force magnitude ‘F’ (left column), and each of the three sub-components: ‘F_z_’ top right, ‘F_x_’ middle right, ‘F_y_’ bottom right columns, for the preferred arm configuration. In each plot the different pushing conditions are presented in different solid color lines: FLAT in dark blue, LCK in light blue, FEX in green, URD in yellow, and ULCK in red. Each individual line represents a single trial. In the total force magnitude plots, two dashed-black lines delimit a shaded grey area representing the considered time window for the analysis of maximum voluntary force performance.

**Figure 18.**
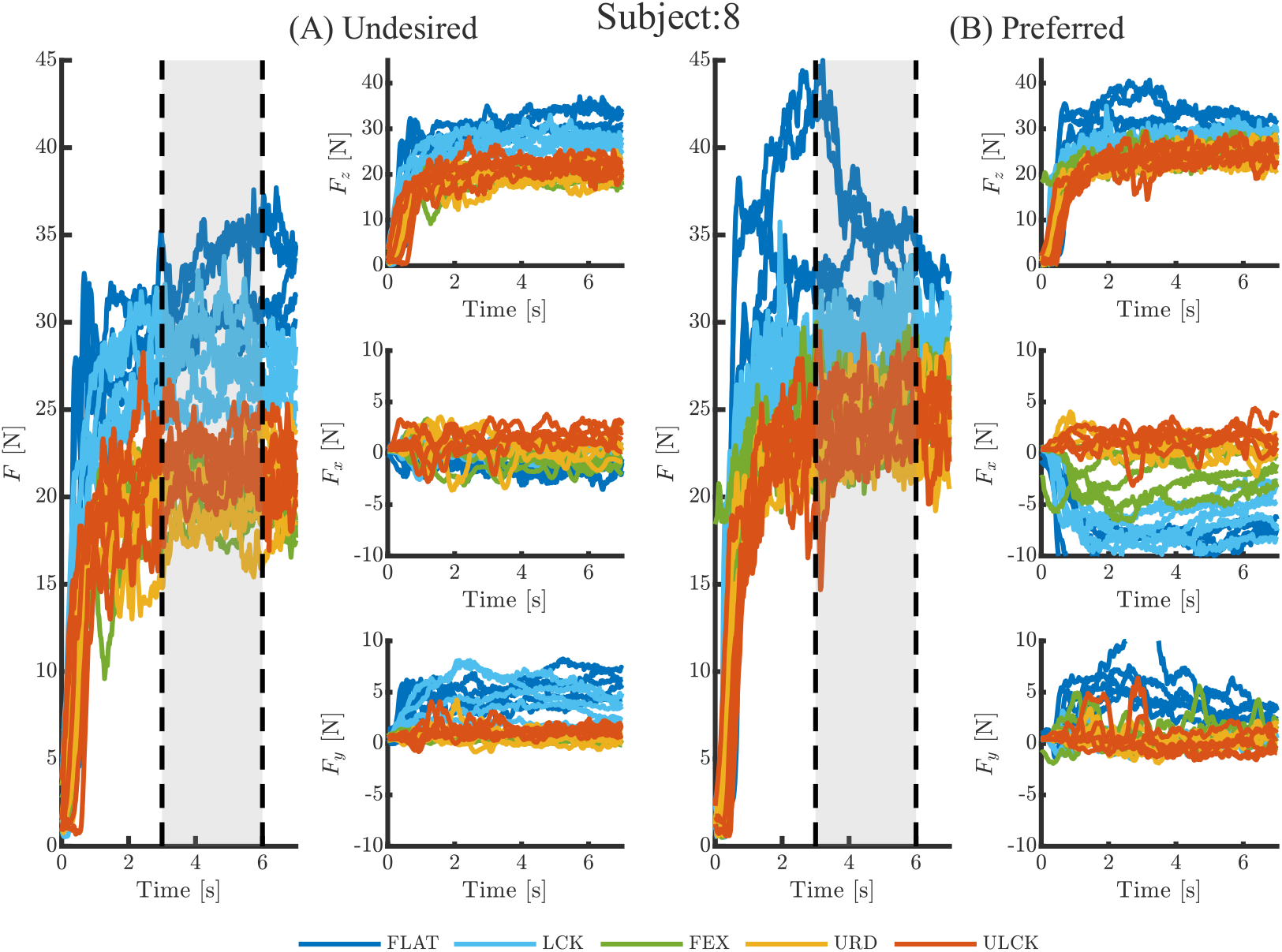
Force temporal trajectories for Subject 8. Panel (A) shows the total force magnitude ‘F’ (left column), and each of the three sub-components: ‘F_z_’ top right, ‘F_x_’ middle right, ‘F_y_’ bottom right columns, for the undesired arm configuration. Panel (B) shows the total force magnitude ‘F’ (left column), and each of the three sub-components: ‘F_z_’ top right, ‘F_x_’ middle right, ‘F_y_’ bottom right columns, for the preferred arm configuration. In each plot the different pushing conditions are presented in different solid color lines: FLAT in dark blue, LCK in light blue, FEX in green, URD in yellow, and ULCK in red. Each individual line represents a single trial. In the total force magnitude plots, two dashed-black lines delimit a shaded grey area representing the considered time window for the analysis of maximum voluntary force performance.

**Figure 19.**
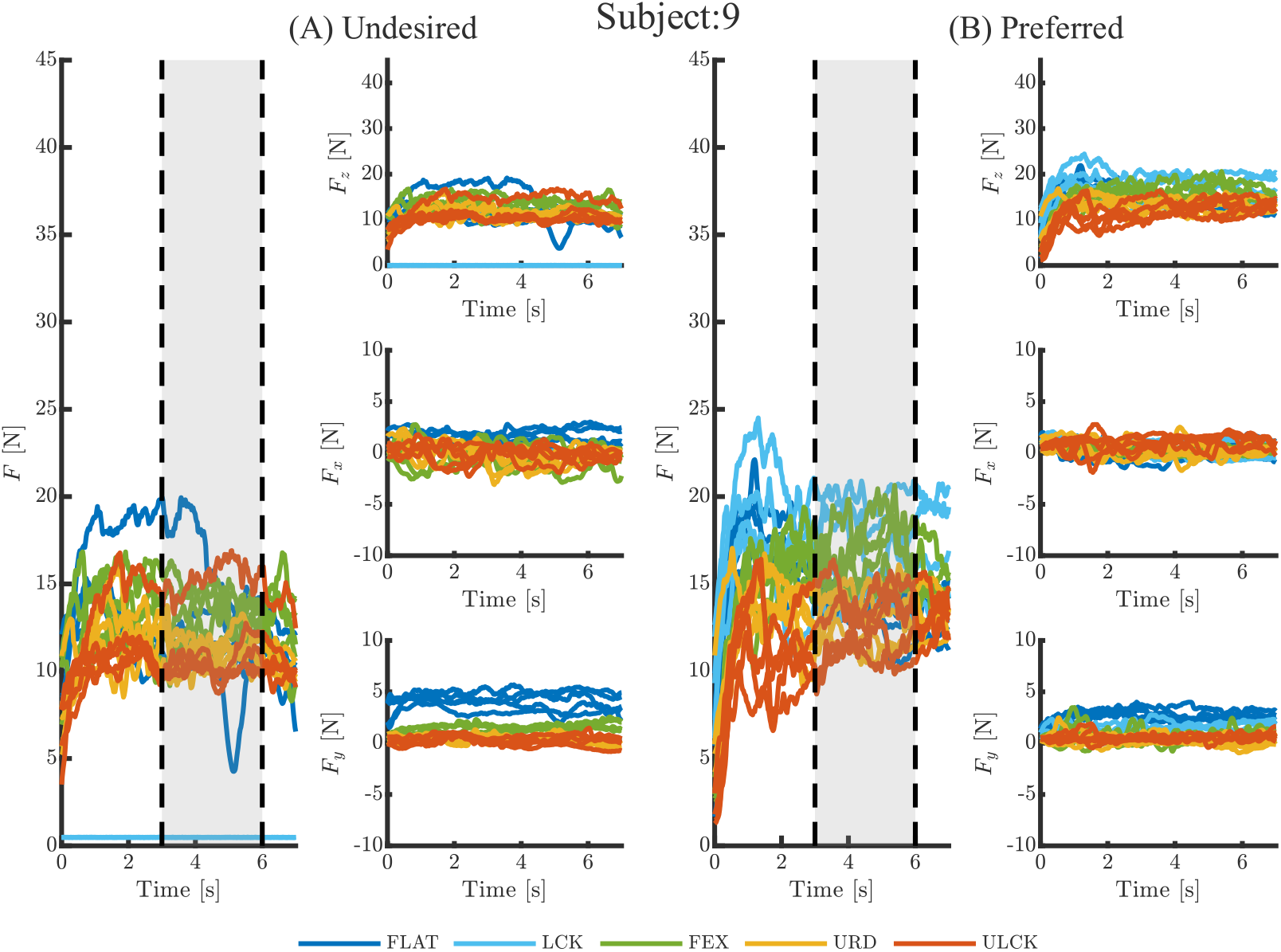
Force temporal trajectories for Subject 9. Panel (A) shows the total force magnitude ‘F’ (left column), and each of the three sub-components: ‘F_z_’ top right, ‘F_x_’ middle right, ‘F_y_’ bottom right columns, for the undesired arm configuration. Panel (B) shows the total force magnitude ‘F’ (left column), and each of the three sub-components: ‘F_z_’ top right, ‘F_x_’ middle right, ‘F_y_’ bottom right columns, for the preferred arm configuration. In each plot the different pushing conditions are presented in different solid color lines: FLAT in dark blue, LCK in light blue, FEX in green, URD in yellow, and ULCK in red. Each individual line represents a single trial. In the total force magnitude plots, two dashed-black lines delimit a shaded grey area representing the considered time window for the analysis of maximum voluntary force performance.

### Statistical Analysis Tables

In the following tables are reported the results of the 2-way repeated measures ANOVA tests performed on each subject to study the effect of arm configuration (undesired, preferred) and task condition (FLAT, LCK, FEX, URD, ULCK) on maximum force generation, as well as the Bonferroni correct post-hoc t-tests.

**Table 2.**
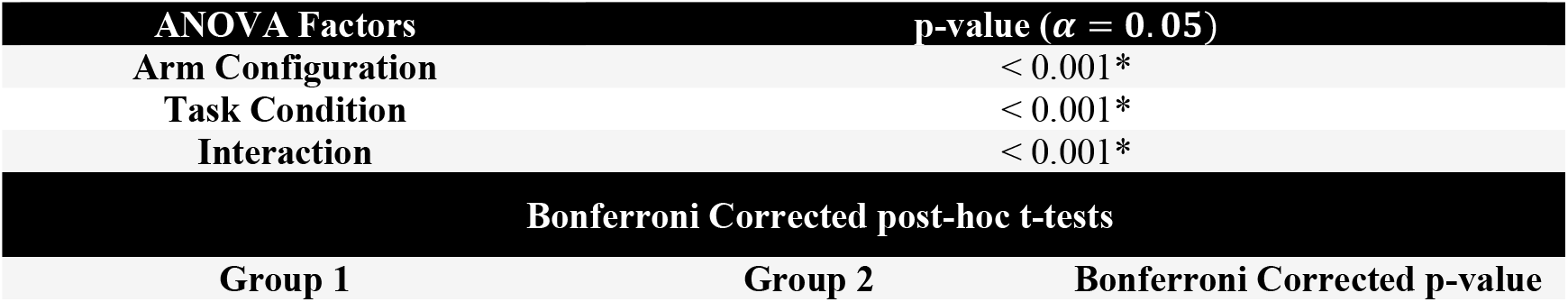
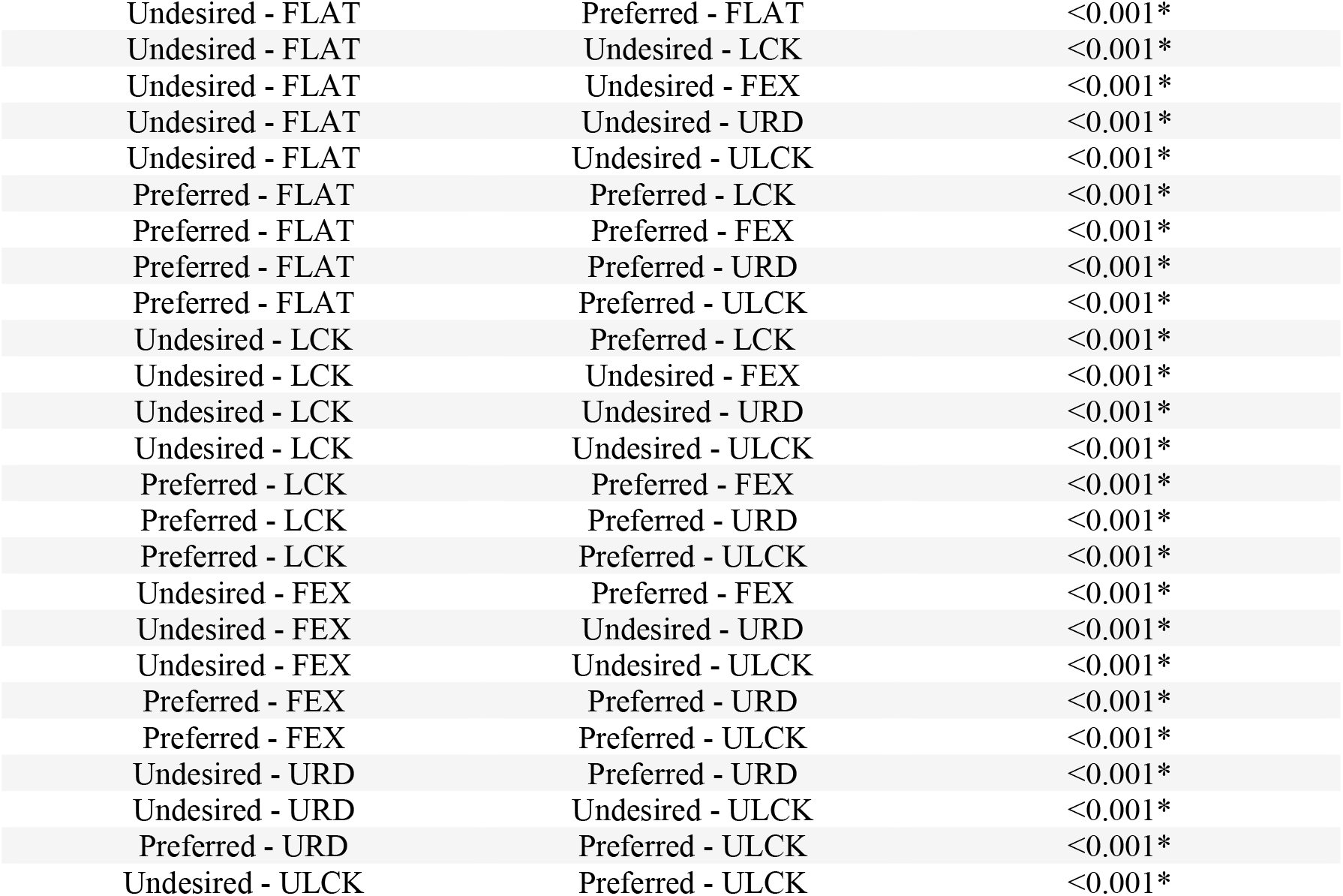
Subject 1 two-way repeated measure ANOVA results to investigate the effect of arm configuration (Undesired, Preferred) and task condition (FLAT, LCK, FEX, URD, ULCK) on the maximum voluntary isometric force. The significant threshold is set at α = 0.05. The post-hoc t-tests are Bonferroni corrected for n=25 comparisons. The asterisk ‘*’ denotes a significant result.

**Table 3.** Subject 2 two-way repeated measure ANOVA results to investigate the effect of arm configuration (Undesired, Preferred) and task condition (FLAT, LCK, FEX, URD, ULCK) on the maximum voluntary isometric force. The significant threshold is set at α = 0.05. The post-hoc t-tests are Bonferroni corrected for n=25 comparisons. The asterisk ‘*’ denotes a significant result.

| ANOVA Factors | | p-value ($\alpha = 0.05$) |
| --- | --- | --- |
| Arm Configuration |  | <0.001* |
| Task Condition |  | <0.001* |
| Interaction |  | <0.001* |
| Bonferroni Corrected post-hoc t-tests |  |  |
| Group 1 | Group 2 | Bonferroni Corrected p-value |
| Undesired - FLAT | Preferred - FLAT | 0.420 |
| Undesired - FLAT | Undesired - LCK | <0.001* |
| Undesired - FLAT | Undesired - FEX | <0.001* |
| Undesired - FLAT | Undesired - URD | <0.001* |
| Undesired - FLAT | Undesired - ULCK | <0.001* |
| Preferred - FLAT | Preferred - LCK | <0.001* |
| Preferred - FLAT | Preferred - FEX | <0.001* |
| Preferred - FLAT | Preferred - URD | <0.001* |
| Preferred - FLAT | Preferred - ULCK | <0.001* |
| Undesired - LCK | Preferred - LCK | <0.001* |
| Undesired - LCK | Undesired - FEX | <0.001* |
| Undesired - LCK | Undesired - URD | <0.001* |
| Undesired - LCK | Undesired - ULCK | <0.001* |
| Preferred - LCK | Preferred - FEX | <0.001* |
| Preferred - LCK | Preferred - URD | <0.001* |
| Preferred - LCK | Preferred - ULCK | <0.001* |
| Undesired - FEX | Preferred - FEX | <0.001* |
| Undesired - FEX | Undesired - URD | <0.001* |
| Undesired - FEX | Undesired - ULCK | <0.001* |
| Preferred - FEX | Preferred - URD | <0.001* |
| Preferred - FEX | Preferred - ULCK | <0.001* |
| Undesired - URD | Preferred - URD | 0.466 |
| Undesired - URD | Undesired - ULCK | <0.001* |
| Preferred - URD | Preferred - ULCK | 1.000 |
| Undesired - ULCK | Preferred - ULCK | <0.001* |

**Table 4.** Subject 3 two-way repeated measure ANOVA results to investigate the effect of arm configuration (Undesired, Preferred) and task condition (FLAT, LCK, FEX, URD, ULCK) on the maximum voluntary isometric force. The significant threshold is set at α = 0.05. The post-hoc t-tests are Bonferroni corrected for n=25 comparisons. The asterisk ‘*’ denotes a significant result.

| ANOVA Factors | | p-value ($\alpha = 0.05$) |
| --- | --- | --- |
| Arm Configuration |  | <0.001* |
| Task Condition |  | <0.001* |
| Interaction |  | <0.001* |
| Bonferroni Corrected post-hoc t-tests |  |  |
| Group 1 | Group 2 | Bonferroni Corrected p-value |
| Undesired - FLAT | Preferred - FLAT | <0.001* |
| Undesired - FLAT | Undesired - LCK | <0.001* |
| Undesired - FLAT | Undesired - FEX | <0.001* |
| Undesired - FLAT | Undesired - URD | <0.001* |
| Undesired - FLAT | Undesired - ULCK | <0.001* |
| Preferred - FLAT | Preferred - LCK | <0.001* |
| Preferred - FLAT | Preferred - FEX | <0.001* |
| Preferred - FLAT | Preferred - URD | <0.001* |
| Preferred - FLAT | Preferred - ULCK | <0.001* |
| Undesired - LCK | Preferred - LCK | <0.001* |
| Undesired - LCK | Undesired - FEX | <0.001* |
| Undesired - LCK | Undesired - URD | <0.001* |
| Undesired - LCK | Undesired - ULCK | <0.001* |
| Preferred - LCK | Preferred - FEX | 1.000 |
| Preferred - LCK | Preferred - URD | <0.001* |
| Preferred - LCK | Preferred - ULCK | <0.001* |
| Undesired - FEX | Preferred - FEX | <0.001* |
| Undesired - FEX | Undesired - URD | <0.001* |
| Undesired - FEX | Undesired - ULCK | <0.001* |
| Preferred - FEX | Preferred - URD | <0.001* |
| Preferred - FEX | Preferred - ULCK | <0.001* |
| Undesired - URD | Preferred - URD | <0.001* |
| Undesired - URD | Undesired - ULCK | <0.001* |
| Preferred - URD | Preferred - ULCK | <0.001* |
| Undesired - ULCK | Preferred - ULCK | <0.001* |

**Table 5.** Subject 4 two-way repeated measure ANOVA results to investigate the effect of arm configuration (Undesired, Preferred) and task condition (FLAT, LCK, FEX, URD, ULCK) on the maximum voluntary isometric force. The significant threshold is set at α = 0.05. The post-hoc t-tests are Bonferroni corrected for n=25 comparisons. The asterisk ‘*’ denotes a significant result.

| ANOVA Factors | | p-value ($\alpha = 0.05$) |
| --- | --- | --- |
| Arm Configuration |  | <0.001* |
| Task Condition |  | <0.001* |
| Interaction |  | <0.001* |
| Bonferroni Corrected post-hoc t-tests |  |  |
| Group 1 | Group 2 | Bonferroni Corrected p-value |
| Undesired - FLAT | Preferred - FLAT | 1.000 |
| Undesired - FLAT | Undesired - LCK | <0.001* |
| Undesired - FLAT | Undesired - FEX | <0.001* |
| Undesired - FLAT | Undesired - URD | <0.001* |
| Undesired - FLAT | Undesired - ULCK | <0.001* |
| Preferred - FLAT | Preferred - LCK | <0.001* |
| Preferred - FLAT | Preferred - FEX | <0.001* |
| Preferred - FLAT | Preferred - URD | <0.001* |
| Preferred - FLAT | Preferred - ULCK | <0.001* |
| Undesired - LCK | Preferred - LCK | <0.001* |
| Undesired - LCK | Undesired - FEX | <0.001* |
| Undesired - LCK | Undesired - URD | <0.001* |
| Undesired - LCK | Undesired - ULCK | <0.001* |
| Preferred - LCK | Preferred - FEX | <0.001* |
| Preferred - LCK | Preferred - URD | <0.001* |
| Preferred - LCK | Preferred - ULCK | <0.001* |
| Undesired - FEX | Preferred - FEX | <0.001* |
| Undesired - FEX | Undesired - URD | <0.001* |
| Undesired - FEX | Undesired - ULCK | <0.001* |
| Preferred - FEX | Preferred - URD | <0.001* |
| Preferred - FEX | Preferred - ULCK | <0.001* |
| Undesired - URD | Preferred - URD | 1.000 |
| Undesired - URD | Undesired - ULCK | <0.001* |
| Preferred - URD | Preferred - ULCK | <0.001* |
| Undesired - ULCK | Preferred - ULCK | 1.000 |

**Table 6.**
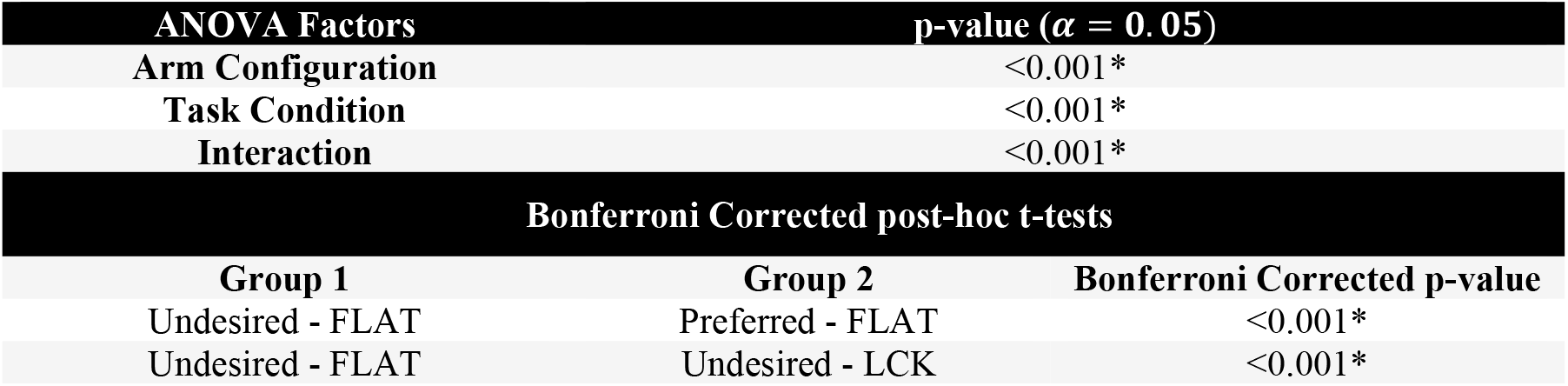
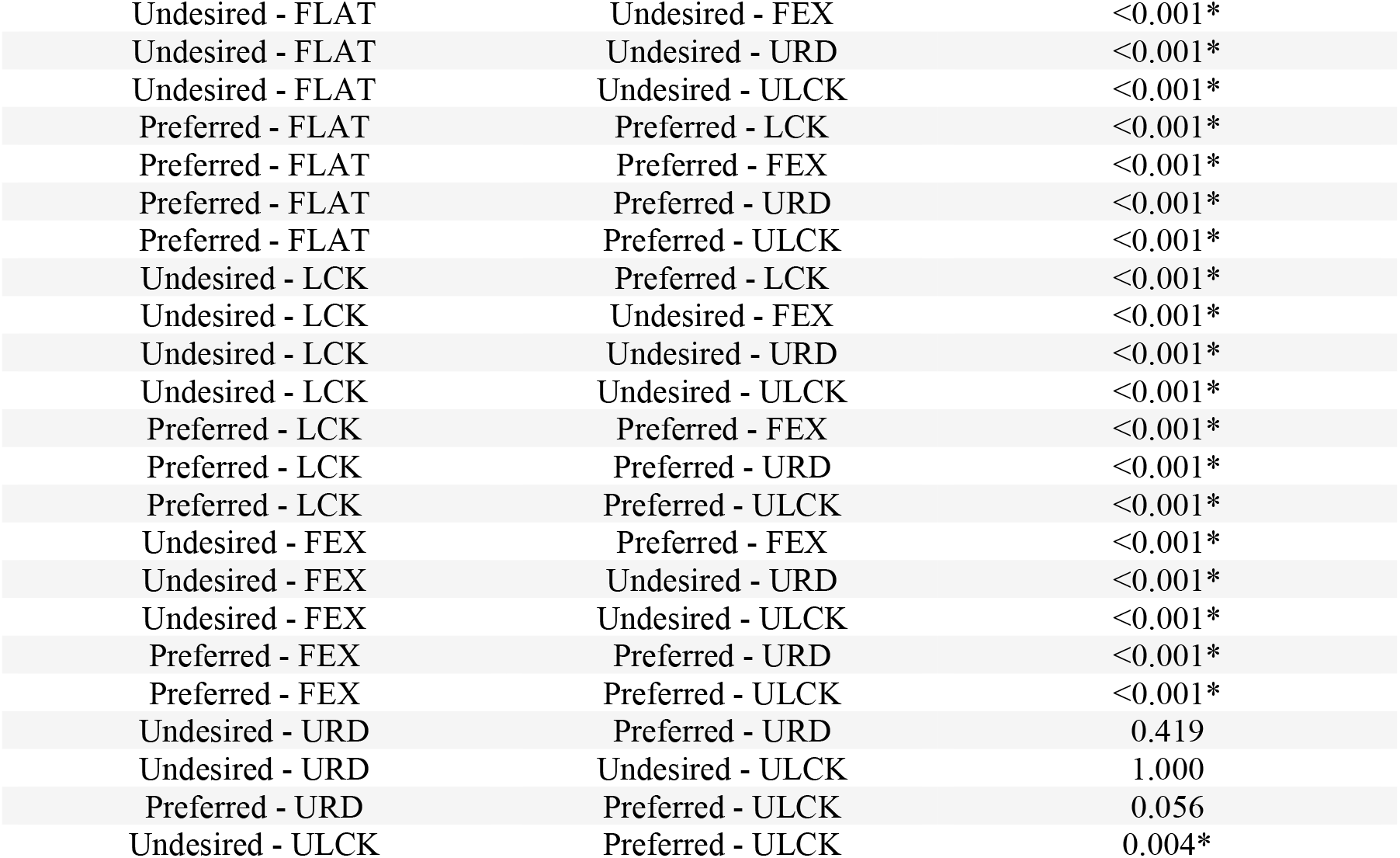
Subject 5 two-way repeated measure ANOVA results to investigate the effect of arm configuration (Undesired, Preferred) and task condition (FLAT, LCK, FEX, URD, ULCK) on the maximum voluntary isometric force. The significant threshold is set at α = 0.05. The post-hoc t-tests are Bonferroni corrected for n=25 comparisons. The asterisk ‘*’ denotes a significant result.

| ANOVA Factors | | p-value ($\alpha = 0.05$) |
| --- | --- | --- |
| Arm Configuration |  | <0.001* |
| Task Condition |  | <0.001* |
| Interaction |  | <0.001* |
| Bonferroni Corrected post-hoc t-tests |  |  |
| Group 1 | Group 2 | Bonferroni Corrected p-value |
| Undesired - FLAT | Preferred - FLAT | <0.001* |
| Undesired - FLAT | Undesired - LCK | <0.001* |
| Undesired - FLAT | Undesired - FEX | <0.001* |
| Undesired - FLAT | Undesired - URD | <0.001* |
| Undesired - FLAT | Undesired - ULCK | <0.001* |
| Preferred - FLAT | Preferred - LCK | <0.001* |
| Preferred - FLAT | Preferred - FEX | <0.001* |
| Preferred - FLAT | Preferred - URD | <0.001* |
| Preferred - FLAT | Preferred - ULCK | <0.001* |
| Undesired - LCK | Preferred - LCK | <0.001* |
| Undesired - LCK | Undesired - FEX | <0.001* |
| Undesired - LCK | Undesired - URD | <0.001* |
| Undesired - LCK | Undesired - ULCK | <0.001* |
| Preferred - LCK | Preferred - FEX | <0.001* |
| Preferred - LCK | Preferred - URD | <0.001* |
| Preferred - LCK | Preferred - ULCK | <0.001* |
| Undesired - FEX | Preferred - FEX | <0.001* |
| Undesired - FEX | Undesired - URD | <0.001* |
| Undesired - FEX | Undesired - ULCK | <0.001* |
| Preferred - FEX | Preferred - URD | <0.001* |
| Preferred - FEX | Preferred - ULCK | <0.001* |
| Undesired - URD | Preferred - URD | 0.419 |
| Undesired - URD | Undesired - ULCK | 1.000 |
| Preferred - URD | Preferred - ULCK | 0.056 |
| Undesired - ULCK | Preferred - ULCK | 0.004* |

**Table 7.** Subject 6 two-way repeated measure ANOVA results to investigate the effect of arm configuration (Undesired, Preferred) and task condition (FLAT, LCK, FEX, URD, ULCK) on the maximum voluntary isometric force. The significant threshold is set at α = 0.05. The post-hoc t-tests are Bonferroni corrected for n=25 comparisons. The asterisk ‘*’ denotes a significant result.

| ANOVA Factors | | p-value ($\alpha = 0.05$) |
| --- | --- | --- |
| Arm Configuration |  | <0.001* |
| Task Condition |  | <0.001* |
| Interaction |  | <0.001* |
| Bonferroni Corrected post-hoc t-tests |  |  |
| Group 1 | Group 2 | Bonferroni Corrected p-value |
| Undesired - FLAT | Preferred - FLAT | <0.001* |
| Undesired - FLAT | Undesired - LCK | <0.001* |
| Undesired - FLAT | Undesired - FEX | <0.001* |
| Undesired - FLAT | Undesired - URD | <0.001* |
| Undesired - FLAT | Undesired - ULCK | <0.001* |
| Preferred - FLAT | Preferred - LCK | <0.001* |
| Preferred - FLAT | Preferred - FEX | <0.001* |
| Preferred - FLAT | Preferred - URD | <0.001* |
| Preferred - FLAT | Preferred - ULCK | <0.001* |
| Undesired - LCK | Preferred - LCK | <0.001* |
| Undesired - LCK | Undesired - FEX | <0.001* |
| Undesired - LCK | Undesired - URD | <0.001* |
| Undesired - LCK | Undesired - ULCK | <0.001* |
| Preferred - LCK | Preferred - FEX | <0.001* |
| Preferred - LCK | Preferred - URD | <0.001* |
| Preferred - LCK | Preferred - ULCK | <0.001* |
| Undesired - FEX | Preferred - FEX | <0.001* |
| Undesired - FEX | Undesired - URD | <0.001* |
| Undesired - FEX | Undesired - ULCK | <0.001* |
| Preferred - FEX | Preferred - URD | <0.001* |
| Preferred - FEX | Preferred - ULCK | <0.001* |
| Undesired - URD | Preferred - URD | <0.001* |
| Undesired - URD | Undesired - ULCK | <0.001* |
| Preferred - URD | Preferred - ULCK | <0.001* |
| Undesired - ULCK | Preferred - ULCK | <0.001* |

**Table 8.** Subject 7 two-way repeated measure ANOVA results to investigate the effect of arm configuration (Undesired, Preferred) and task condition (FLAT, LCK, FEX, URD, ULCK) on the maximum voluntary isometric force. The significant threshold is set at α = 0.05. The post-hoc t-tests are Bonferroni corrected for n=25 comparisons. The asterisk ‘*’ denotes a significant result.

| ANOVA Factors | | p-value ($\alpha = 0.05$) |
| --- | --- | --- |
| Arm Configuration |  | <0.001* |
| Task Condition |  | <0.001* |
| Interaction |  | <0.001* |
| Bonferroni Corrected post-hoc t-tests |  |  |
| Group 1 | Group 2 | Bonferroni Corrected p-value |
| Undesired - FLAT | Preferred - FLAT | <0.001* |
| Undesired - FLAT | Undesired - LCK | <0.001* |
| Undesired - FLAT | Undesired - FEX | <0.001* |
| Undesired - FLAT | Undesired - URD | <0.001* |
| Undesired - FLAT | Undesired - ULCK | <0.001* |
| Preferred - FLAT | Preferred - LCK | <0.001* |
| Preferred - FLAT | Preferred - FEX | <0.001* |
| Preferred - FLAT | Preferred - URD | <0.001* |
| Preferred - FLAT | Preferred - ULCK | <0.001* |
| Undesired - LCK | Preferred - LCK | <0.001* |
| Undesired - LCK | Undesired - FEX | <0.001* |
| Undesired - LCK | Undesired - URD | <0.001* |
| Undesired - LCK | Undesired - ULCK | <0.001* |
| Preferred - LCK | Preferred - FEX | <0.001* |
| Preferred - LCK | Preferred - URD | <0.001* |
| Preferred - LCK | Preferred - ULCK | <0.001* |
| Undesired - FEX | Preferred - FEX | <0.001* |
| Undesired - FEX | Undesired - URD | 0.096 |
| Undesired - FEX | Undesired - ULCK | <0.001* |
| Preferred - FEX | Preferred - URD | <0.001* |
| Preferred - FEX | Preferred - ULCK | <0.001* |
| Undesired - URD | Preferred - URD | <0.001* |
| Undesired - URD | Undesired - ULCK | <0.001* |
| Preferred - URD | Preferred - ULCK | <0.001* |
| Undesired - ULCK | Preferred - ULCK | <0.001* |

**Table 9.** Subject 8 two-way repeated measure ANOVA results to investigate the effect of arm configuration (Undesired, Preferred) and task condition (FLAT, LCK, FEX, URD, ULCK) on the maximum voluntary isometric force. The significant threshold is set at α = 0.05. The post-hoc t-tests are Bonferroni corrected for n=25 comparisons. The asterisk ‘*’ denotes a significant result.

| ANOVA Factors | | p-value ($\alpha = 0.05$) |
| --- | --- | --- |
| Arm Configuration |  | <0.001* |
| Task Condition |  | <0.001* |
| Interaction |  | <0.001* |
| Bonferroni Corrected post-hoc t-tests |  |  |
| Group 1 | Group 2 | Bonferroni Corrected p-value |
| Undesired - FLAT | Preferred - FLAT | <0.001* |
| Undesired - FLAT | Undesired - LCK | <0.001* |
| Undesired - FLAT | Undesired - FEX | <0.001* |
| Undesired - FLAT | Undesired - URD | <0.001* |
| Undesired - FLAT | Undesired - ULCK | <0.001* |
| Preferred - FLAT | Preferred - LCK | <0.001* |
| Preferred - FLAT | Preferred - FEX | <0.001* |
| Preferred - FLAT | Preferred - URD | <0.001* |
| Preferred - FLAT | Preferred - ULCK | <0.001* |
| Undesired - LCK | Preferred - LCK | <0.001* |
| Undesired - LCK | Undesired - FEX | <0.001* |
| Undesired - LCK | Undesired - URD | <0.001* |
| Undesired - LCK | Undesired - ULCK | <0.001* |
| Preferred - LCK | Preferred - FEX | <0.001* |
| Preferred - LCK | Preferred - URD | <0.001* |
| Preferred - LCK | Preferred - ULCK | <0.001* |
| Undesired - FEX | Preferred - FEX | <0.001* |
| Undesired - FEX | Undesired - URD | <0.001* |
| Undesired - FEX | Undesired - ULCK | <0.001* |
| Preferred - FEX | Preferred - URD | <0.001* |
| Preferred - FEX | Preferred - ULCK | <0.001* |
| Undesired - URD | Preferred - URD | <0.001* |
| Undesired - URD | Undesired - ULCK | <0.001* |
| Preferred - URD | Preferred - ULCK | <0.001* |
| Undesired - ULCK | Preferred - ULCK | <0.001* |

**Table 10.** Subject 9 two-way repeated measure ANOVA results to investigate the effect of arm configuration (Undesired, Preferred) and task condition (FLAT, LCK, FEX, URD, ULCK) on the maximum voluntary isometric force. The significant threshold is set at α = 0.05. The post-hoc t-tests are Bonferroni corrected for n=25 comparisons. The asterisk ‘*’ denotes a significant result.

| ANOVA Factors | | p-value ($\alpha = 0.05$) |
| --- | --- | --- |
| Arm Configuration |  | <0.001* |
| Task Condition |  | <0.001* |
| Interaction |  | <0.001* |
| Bonferroni Corrected post-hoc t-tests |  |  |
| Group 1 | Group 2 | Bonferroni Corrected p-value |
| Undesired - FLAT | Preferred - FLAT | <0.001* |
| Undesired - FLAT | Undesired - LCK | <0.001* |
| Undesired - FLAT | Undesired - FEX | <0.001* |
| Undesired - FLAT | Undesired - URD | <0.001* |
| Undesired - FLAT | Undesired - ULCK | <0.001* |
| Preferred - FLAT | Preferred - LCK | <0.001* |
| Preferred - FLAT | Preferred - FEX | <0.001* |
| Preferred - FLAT | Preferred - URD | <0.001* |
| Preferred - FLAT | Preferred - ULCK | <0.001* |
| Undesired - LCK | Preferred - LCK | <0.001* |
| Undesired - LCK | Undesired - FEX | <0.001* |
| Undesired - LCK | Undesired - URD | <0.001* |
| Undesired - LCK | Undesired - ULCK | <0.001* |
| Preferred - LCK | Preferred - FEX | <0.001* |
| Preferred - LCK | Preferred - URD | <0.001* |
| Preferred - LCK | Preferred - ULCK | <0.001* |
| Undesired - FEX | Preferred - FEX | <0.001* |
| Undesired - FEX | Undesired - URD | <0.001* |
| Undesired - FEX | Undesired - ULCK | <0.001* |
| Preferred - FEX | Preferred - URD | <0.001* |
| Preferred - FEX | Preferred - ULCK | <0.001* |
| Undesired - URD | Preferred - URD | <0.001* |
| Undesired - URD | Undesired - ULCK | 1.000 |
| Preferred - URD | Preferred - ULCK | <0.001* |
| Undesired - ULCK | Preferred - ULCK | <0.001* |

## Footnotes

1 In this work we always refer to mechanical impedance, therefore we frequently omit the ‘mechanical’ prefix.

2 We leave the readers the pleasure to derive this equation. The solution is straightforward once you observe that the new lever arm ‘*h*’ of the pushing force ‘*F*’ is equal to *h* = *l* sin(Δ*θ* + Δ*α*) ≈ *l*(Δ*θ* + Δ*α*).

3 The chosen convention for the orientation of the coordinate reference frame replicates the same one adopted by the force/torque sensor during the experiment.

4 Slight variations of the window size were considered but no particular differences were observed.

